# Profiling of extracellular small RNAs highlights a strong bias towards non-vesicular secretion

**DOI:** 10.1101/2020.12.01.406207

**Authors:** Helena Sork, Mariana Conceicao, Giulia Corso, Joel Nordin, Yi Xin Fiona Lee, Kaarel Krjutskov, Jakub Orzechowski Westholm, Pieter Vader, Marie Pauwels, Roosmarijn Vandenbroucke, Matthew JA Wood, Samir EL Andaloussi, Imre Mäger

## Abstract

Extracellular environment consists of a plethora of different molecules, including extracellular miRNA that can be secreted in association with extracellular vesicles (EVs) or soluble protein complexes (non-EVs). Yet, it is generally accepted that most of the biological activity is attributed to EV-associated miRNAs. The capability of EVs to transport cargoes has attracted much interest towards developing EVs as therapeutic short RNA carriers by using endogenous loading strategies for miRNA enrichment. Here, by overexpressing miRNA and shRNA sequences of interest in source cells and using size exclusion liquid chromatography (SEC) to separate the cellular secretome into EV and non-EV fractions, we saw that strikingly, <2% of all secreted overexpressed miRNA were found in association with EVs. To see whether the prominent non-EV miRNA secretion also holds true at the basal expression level of native miRNA transcripts, both fractions were further analysed by small RNA sequencing. This revealed a global correlation of EV and non-EV miRNA abundance to that of their parent cells and showed an enrichment only for miRNAs with a relatively low cellular expression level. Further quantification showed that similarly to the transient overexpression context, an outstanding 96.2-99.9% of total secreted miRNA at its basal level was secreted to the non-EV fraction. Yet, even though EVs contain only a fraction of secreted miRNAs, these molecules were found stable at 37°C in serum-containing environment, indicating that if sufficient miRNA loading to EVs is achieved, EVs can remain miRNA delivery-competent for a prolonged period of time. This study suggests that the passive endogenous EV loading strategy can be a relatively wasteful way of loading miRNA to EVs and active miRNA loading approaches are needed for developing advanced EV miRNA therapies in the future.

## INTRODUCTION

The constant drive to find novel carriers of RNA therapeutics has witnessed a shift from artificial delivery systems to nature-inspired biomolecule carriers such as extracellular vesicles (EVs). Owing to their pivotal role in cell to cell communication as well as the inherent ability to deliver bioactive lipids, proteins and nucleic, EVs hold a spotlight position in the research of advanced therapeutic carriers.

EVs contain a wide range of nucleic acid molecules and though stretches of up to several kilobases in size have been detected^1^, the majority of the encapsulated material does not exceed 200 nucleotides^2,3^. Therefore, most studies focus on loading EVs with therapeutic miRNAs and other short miRNA-like RNAs such as siRNAs and shRNAs^4–6^. In order to load EVs with RNA, exogenous loading as well as endogenous overexpression of therapeutic genes has been utilized, the latter often being preferred^7–9^. Nevertheless, despite significant effort, the enrichment of EVs with the nucleic acid of interest still remains challenging.

It is evident from previous studies that most of the extracellular nucleic acids in plasma is not associated with vesicular structures, being either bound to protein complexes or lipoprotein particles^10,11^. Yet, it is generally accepted that practically all the biological activity is rather conveyed by EVs than other circulating nucleic acid carriers. The presence of a distinct miRNA per EV has been estimated to approximately 1 in 100, or even 1 in 10,000^12–15^ being largely insufficient to ensure solid therapeutic delivery^16^. This underscores the urgency to understand miRNA (and more broadly short RNA) accumulation in EVs and non-EVs and could provide important background information on which strategies to explore for enriching the desired short RNAs to EVs. Different loading strategies have already been employed^17^, but frequently without considering the context of global miRNA secretion.

To be able to harness the potential of EV-encapsulated nucleic acid delivery for biomedical applications, we must increase our understanding of the composition and abundance of short RNAs in different fractions of the extracellular secretome. In this study, we separated cell culture supernatants into EVs and non-vesicular material using SEC and characterized basally expressed and overexpressed miRNA levels in both fractions, in order to better understand important aspects of secretory miRNA sorting.

## RESULTS

It is known that cells constantly release miRNA to the extracellular space. A fraction of secreted miRNA is encapsulated in EVs, yet according to previous reports^10,11^, a large amount can be detected outside vesicular structures. To be able to separate the secretome into vesicular (EVs) and non-vesicular fractions (non-EVs) this study has employed size-exclusion chromatography (SEC) (Figure 1A). We have previously validated SEC workflow for its suitability to separate EVs from the non-vesicular secreted material and found that the total yield of the entire EV isolation workflow was high (> 70%)^18^. Furthermore, the isolated EVs had high purity as EV protein markers were detected only in the SEC eluate, corresponding to the void volume peak (i.e. the EV peak) and not in fractions that eluted later (i.e. the non-EV peak)^18^.

**Figure 1.**
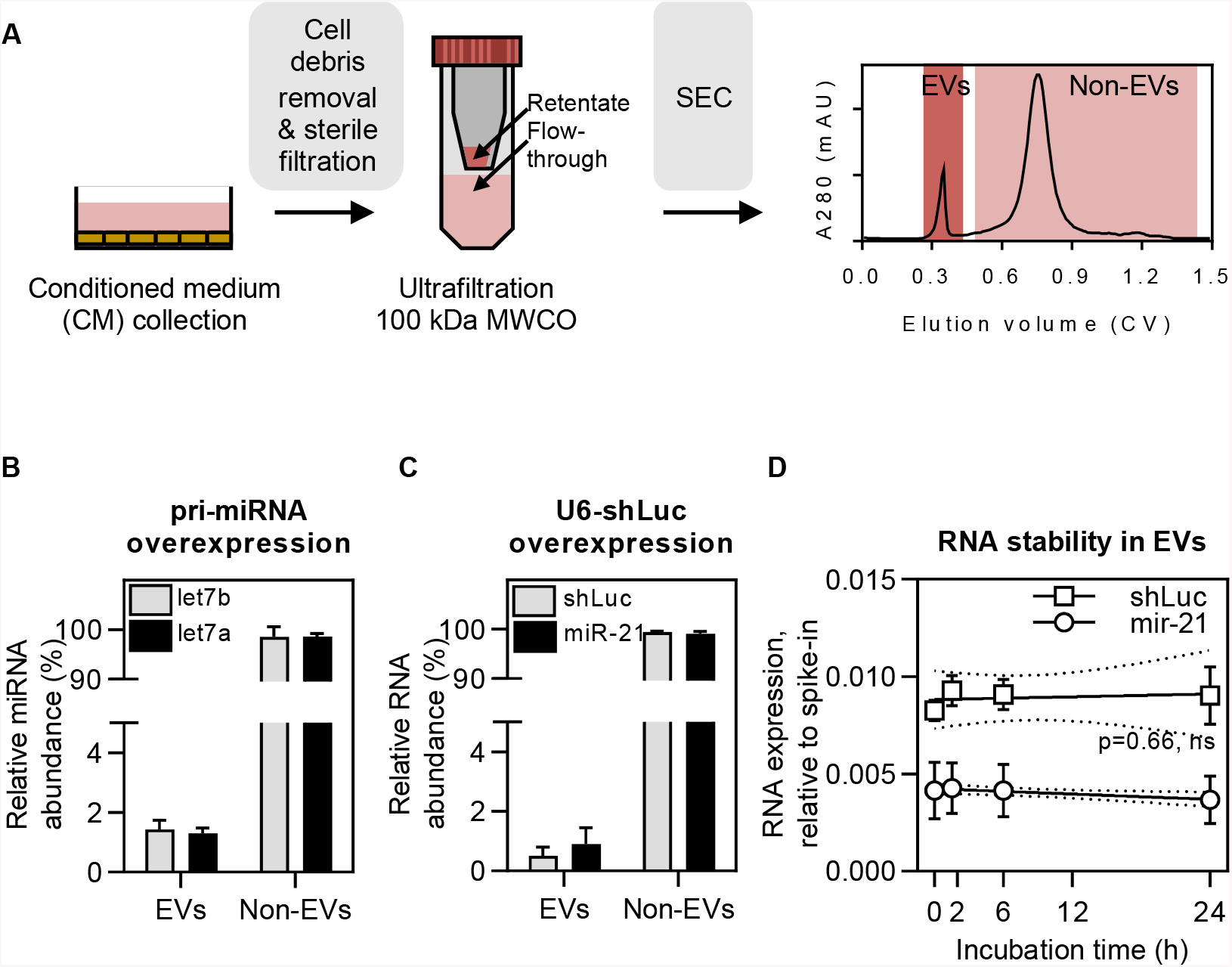
EV isolation workflow and the secretory distribution and stability of overexpressed miRNAs. **(A)** HEK293T (HEK) cells and hTERT-MSCs (MSCs) were seeded to 15-cm tissue culture plates. For producing conditioned medium (CM), cells were incubated in Opti-MEM for 48 hours. For CM fractionation, it was first cleared from cell debris and large particles using centrifugation and sterile filtration, concentrated by ultrafiltration and thereafter fractionated into EV and Non-EV fractions using SEC with Sephacryl S-400 resin. **(B, C)** miRNA sequences were transiently overexpressed in cells using pri-miRNA or U6-shRNA expression plasmids, followed by miRNA quantification using RT-qPCR. Strikingly >98% of overexpressed miRNAs were found to be secreted to the Non-EV fraction. Nevertheless, despite that only a minor fraction of miRNA was found in EVs, these miRNAs showed high serum stability for up to 24 hours **(D)**.

In the present study, SEC-purified EVs were characterised by Western blot, Nanoparticle Tracking Analysis (NTA) and transmission electron microscopy (TEM) (Suppl. Fig. 1). Further analysis of eluted SEC fractions confirmed that high molecular weight particles, as detected by NTA, eluted all in the void volume peak (Suppl. Fig. 2A, B). Also, as expected, most of the protein and RNA was found to elute in the non-EV area of the chromatogram with the latter being susceptible to RNAseA treatment (Suppl.Fig. 2C, D). Additional control experiments with 100 kDa MWCO ultrafiltration spin filters showed that all detectable particles remained in the retentate while most of the miRNA was found in the filtrate (Suppl. Fig. 2E, F), corroborating with the RNA elution profile of the SEC chromatogram. Due to the widely recognised concerns on bovine serum derived miRNA contamination (even when EV-depleted serum is used)^19,20^, the conditioned medium was prepared using serum free Opti-MEM, being suitable for EV production as shown previously^21,22^.

### Pri-miRNA and U6-shRNA overexpression leads to RNA secretion in the non-EV fraction

Many studies focusing on developing EVs as therapeutic RNA carriers aim to enrich miRNA in EVs by overexpressing the respective transcripts in EV producing cells. Yet, little is known about miRNA secretion patterns when using such strategy. To investigate to which extent miRNAs of interest get released to the extracellular space, we transiently overexpressed pri-let-7a and pri-let-7b in HEK cells and quantified the respective mature miRNA levels in the EV and non-EV portion of the secretome. This analysis revealed that the majority of overexpressed let-7a and let-7b mature transcripts were secreted in the non-EV fraction, whereas the EV fraction contained < 2% of the respective miRNAs (Figure 1B). This suggests that while miRNA overexpression in source cells does lead to its increased level in EVs, the majority of extracellular transcripts are predominantly found in the non-EV fraction.

Of note, the pri-miRNA expression vectors used above were driven by the CMV promoter, which is a Pol II promoter and requires full processing of the overexpressed pri-miRNA transcripts by respective miRNA processing machinery (DROSHA, DICER1, etc.). However, in many cases when aiming to load EVs with small RNA, the transcription is initiated using the U6 (Pol III) promoter. This promoter facilitates highly efficient RNA transcription with well-defined transcription start and termination sites, being extensively used for expressing short hairpin RNA (shRNA) for siRNA-like gene silencing purposes. Having this in mind we also overexpressed a shRNA transcript (shRNA targeted against Firefly luciferase, shLuc) in HEK cells to measure its effect on shRNA secretion in EVs. Interestingly, even though the U6-promoter-driven shRNA expression level was significantly higher than the miRNA expression level using the CMV-driven pri-miRNA expression vectors, we observed that the shRNA secretion profile remained unchanged. Namely, <2% of all secreted shLuc was found in the EVs, where similarly to naturally secreted miR-21 transcripts it remained stable in serum-containing environment for 24 h (Figure 1C, D). This indicates that even though only a minority of miRNA is secreted with EVs, it remains stable over a long period of time and may thus be biologically active when delivered to recipient cells by EVs.

### Small RNA secretome is predominantly composed of miRNA and piRNA-like sequences

After observing that the overexpressed miRNAs are predominantly secreted via a non-vesicular route, we were curious to investigate whether the same trend applies to miRNAs on their basal expression level. Hence, we sequenced RNA from cells, SEC-derived EV and non-EV fractions from HEK and MSC cell lines and focused on the analysis of reads between 17-35 nucleotides in length. This restriction was applied in order to not only cover full-length miRNAs, but to also include other small RNAs (sRNA) and check for the presence of larger RNA fragments. The latter have been found in EVs by us and others even when focusing on short reads (Figure 2A)^23–25^.

**Figure 2.**
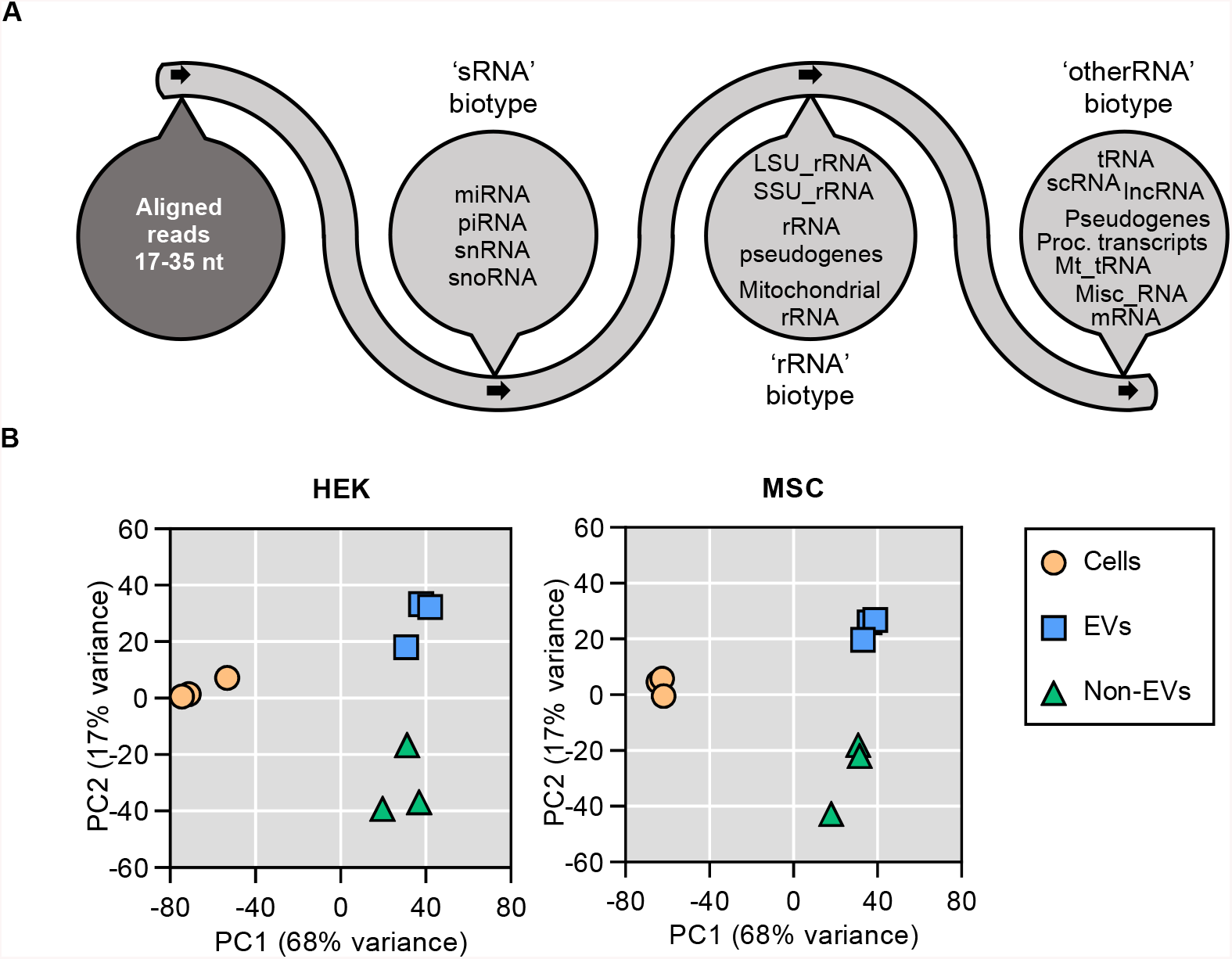
Sequencing workflow and Principle Component Analysis of all uniquely annotated reads. **(A)** RNA was extracted from EV and Non-EV fractions and subjected to sequencing. RNA sequences with a read length of 17–35 nt were aligned to the human genome and annotated in a stepwise manner first to the sRNA biotype (miRNA, piRNA, snRNA, snoRNA), rRNA biotype (LSU_rRNA, SSU_rRNA, rRNA pseudogenes, mitochondrial rRNA), and otherRNA biotype (tNRA, scRNA, lncRNA, pseudogenes, processed transcripts, mt_tRNA, misc_RNA, mRNA). **(B)**Principle Component Analysis of all uniquely annotated reads of both HEK (left panel) and MSC (right panel) samples showed clear cluster separation of cell, EV and Non-EV samples. Looking at the principle component 1 (PC1, explaining 68% of variance) the secretory fractions of both samples are more similar to each other than to parental cells.

Principal component analysis (PCA) showed a clear cluster separation between cell, EV and non-EV samples, indicating to significant differences in their RNA composition (Figure 2B). Upon detailed analysis of the sRNA secretome, we found that miRNA and piRNA-like sequences together constituted the majority of sRNA biotype reads (Figure 3A, B) and depending on the secretory fraction and cell type, covered between 88-98% of sRNA sequences. When comparing with piRNA and miRNA levels in parental cells, piRNA reads were enriched in both EV and non-EV samples, whereas miRNAs were not (Suppl. Fig. 3A, B). Due to sequence similarities and conflicting data in piRNA databases, the sequences which aligned to piRNA loci were also checked for the presence of piRNA features (such as read length distribution and the presence of 5’ uracil) and potential overlap with tRNA annotations (Suppl. Fig. 3C–H). The highest level of tRNA-overlapping sequences was found in non-EV samples, yet even when the potentially tRNA-derived sequences were removed from the analysis, the trend of EVs being enriched in piRNA sequences (compared to their parent cells) remained largely unchanged (Suppl. Fig. 3G, H).

**Figure 3.**
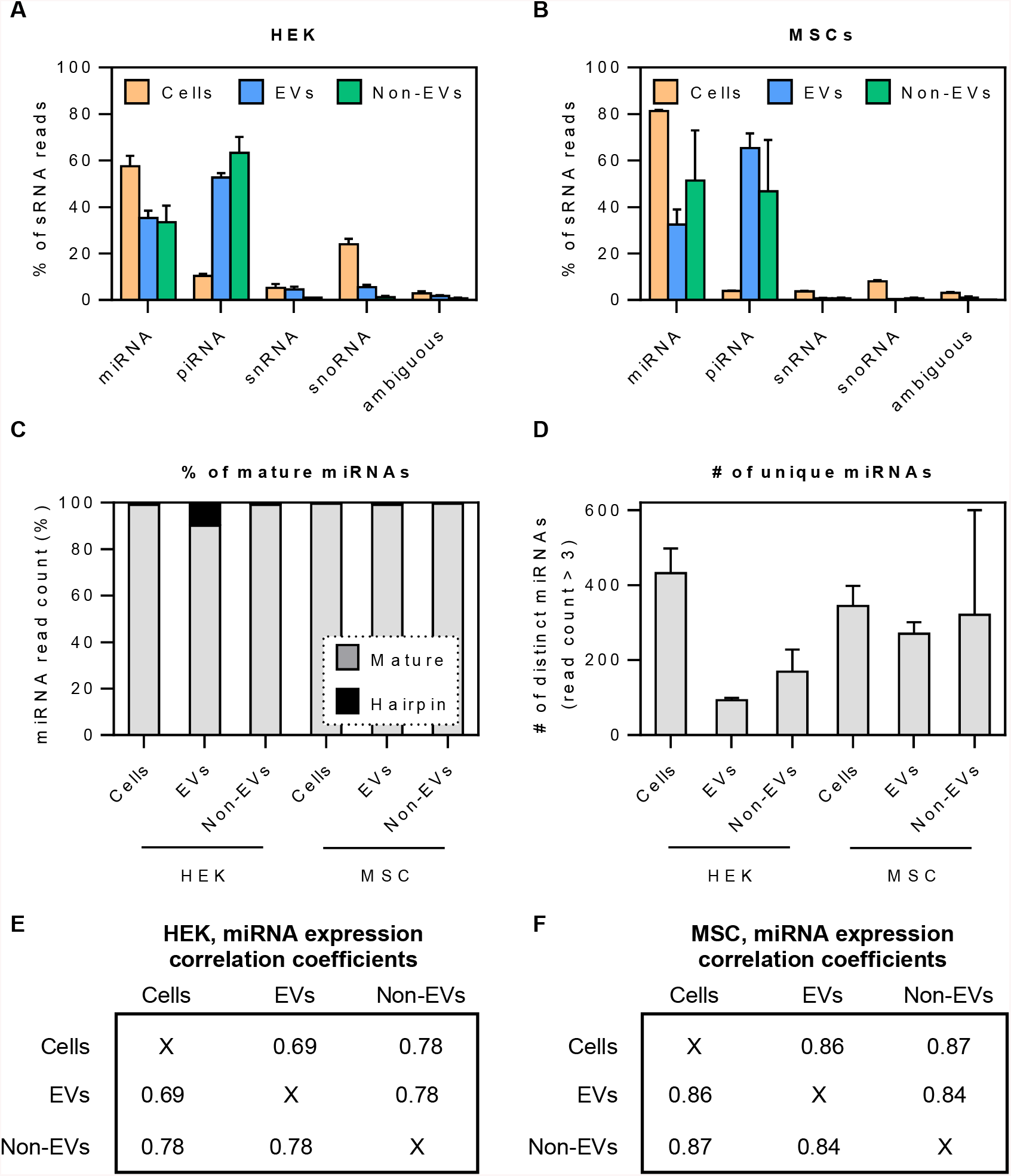
sRNA composition, characteristics and correlation in source cells and secretory fractions. **(A, B)** Within the sRNA biotype, both for HEK and MSC samples, most of the reads comprised of miRNA and piRNA sequences. While cellular RNA was rich in miRNA, EV and Non-EV samples were dominated by sequences aligning to piRNA genes. **(C, D)** The vast majority of detected miRNA sequences were fully processed and the number of distinct miRNAs was generally higher in cell samples than in EVs, particularly for HEK samples. **(E, F)**The correlation of miRNA expression between cellular, EV and Non-EV samples was high, ranging from Pearson R^2^=0.69–0.87.

Overall, it must be noted that the proportion of sRNA biotype sequences in the RNA secretome was relatively low (<20% of total reads within the 17-35 nt read length range). The majority of the annotated sequences were ‘rRNA’ and ‘other RNA’ fragments (Suppl. Fig. 4), the latter mainly covering fragments of tRNA- and protein coding sequences (Suppl. Fig. 5). Secreted miscellaneous RNA in both cell types consisted primarily of Y RNA sequences. Yet, small cytoplasmic RNAs and vault RNAs, which were prominently observed in EVs, remained practically undetected in the non-EV secretome (Suppl. Fig. 5).

### The bulk of secretory miRNAs reflect the content of the source cells

After observing that miRNA and piRNA were the most abundant sRNA species both in EVs and non-EVs, we next decided to analyse whether there are any differences in miRNA secretion between the different fractions. We observed that nearly all miRNA reads fell in the range of 20-24 nucleotides in length and represented mature miRNA sequences (Figure 3C). Interestingly, the number of distinct miRNAs detected in the HEK secretome was lower than in HEK cells, whereas in MSC samples the miRNA numbers were relatively similar in cells, EVs as well as non-EVs (Figure 3D).

The analysis of expression levels of individual miRNAs revealed a good correlation between the EV and non-EV samples (Pearson R^2^=0.78 and R^2^=0.84 for HEK and MSC samples, respectively; Figure 3E, F). When comparing each of the secretory fractions to cells, miRNA expression in both EVs and non-EV fractions correlated highly to parent MSCs (R^2^=0.86 and R^2^=0.87, respectively). The same was seen for the HEK non-EV sample (R^2^=0.78), yet miRNAs in HEK EVs showed a somewhat lower correspondence to their source cells (R^2^=0.69). Nevertheless, these generally high correlation coefficients indicate that, for most miRNAs, a significant contributor to their secretory level is their expression in the cells, suggesting an overall stochastic secretion mechanism.

### Enriched secretory miRNAs have low expression levels in cells

Even though miRNA expression was highly correlated between cells, EVs and non-EVs, there were several individual miRNA sequences that seemed to be enriched in different fractions of the secretome. To confirm that, we performed differential expression (DE) analysis of mature miRNA levels in both cell types by comparing miRNA expression between EVs and non-EVs as well as against their background level in their source cells. As a general observation, miRNAs that were enriched in EVs and in the non-EV fraction displayed low cellular expression level (Figure 4A, B). In HEK-derived samples, a total of 460 distinct miRNAs were detected, out of which, as compared to their source cells, 38 and 31 miRNAs were differentially expressed in EV and non-EV samples, respectively (Figure 4A; Suppl. Table 1). Out of 516 mature miRNAs that were included in the analysis for MSC-derived samples, we detected 78 and 83 DE miRNAs in EV and non-EV samples as compared to source cells (p_adj_ < 0.05, log2 fold change ≥ 1.5; Figure 4B; Suppl. Table 2). As exemplified by the heatmaps and hierarchical clustering analysis, miRNAs with decreased secretion level clustered together for both EV *versus* cells and non-EV *versus* cells comparisons (Figure 4C-F, bright blue branches). This suggests that certain high-expressing miRNAs are likely to be retained in cells via an unknown mechanism.

**Figure 4.**
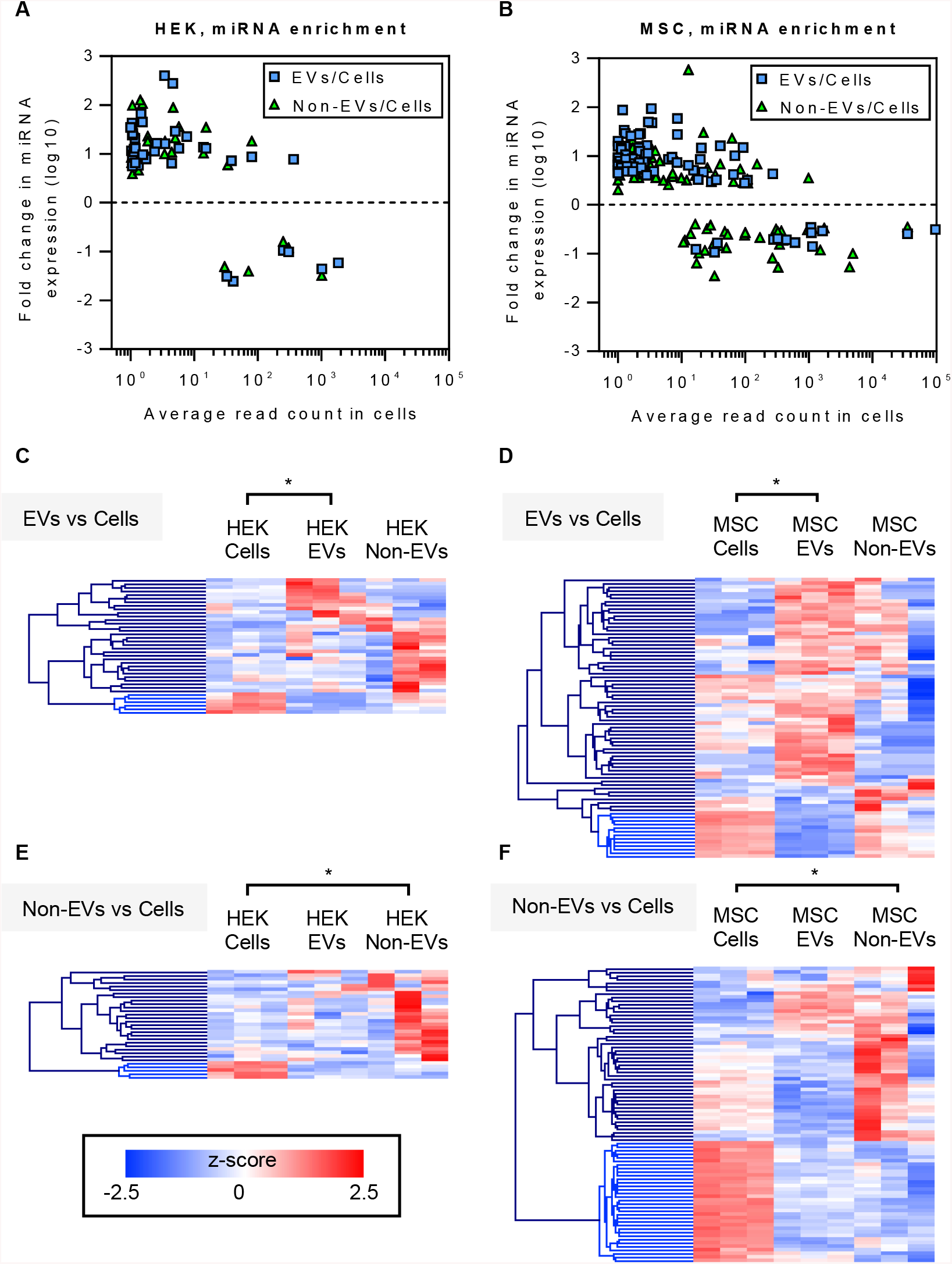
The enrichment pattern of secretory miRNAs as compared to source cells. **(A, B)** miRNA relative enrichment analysis revealed that those miRNAs that were enriched in EVs and Non-EVs had a low average read count in cells. This is true both in case of HEK as well as MSC samples. **(C–F)** Heat maps of hierarchical clustering analysis further suggested that certain miRNAs are restricted from secretion, as miRNAs enriched in cells were reduced both in the EV and Non-EV fractions. Also, the enrichment of miRNAs in EVs was generally not mirrored in the Non-EV fraction.

### The bulk of extracellular miRNAs are secreted via non-EV route

From the sequencing results we could conclude that the enrichment of short RNA sequences at their basal level into EVs can mainly be seen for lowly expressed sequences. To validate these findings with an alternative methodology, we focused on miRNAs with a wide basal expression range (and hence potentially different secretion enrichment characteristics). We selected five miRNAs from our data set (miR-100, miR-210, miR-16, miR-21 and miR-186) and quantified their secretion level by RT-qPCR. As compared to source cells, the level of miR-100 and miR-21 was higher in the extracellular space for HEK cells and lower for MSCs (Suppl. Table 1–2), whereas no significant changes were detected in secreted miR-210, miR-16 and miR-186 (Suppl. Table 1–2). Nevertheless, regardless of the basal expression and secretory enrichment level of tested miRNAs, similarly to overexpression experiments, only a very small proportion of total secreted miRNAs was found associated with EVs. More specifically, only 0.1–0.4% and 0.9–3.8% of miRNA sequences were detected in EVs of HEK- and MSC-derived conditioned medium, respectively (Figure 5A, B). Thus, a striking 96.2–99.9% of extracellular miRNAs were secreted in association with non-EV carriers, underpinning the need for active strategies to enforce vesicular miRNA loading.

**Figure 5.**
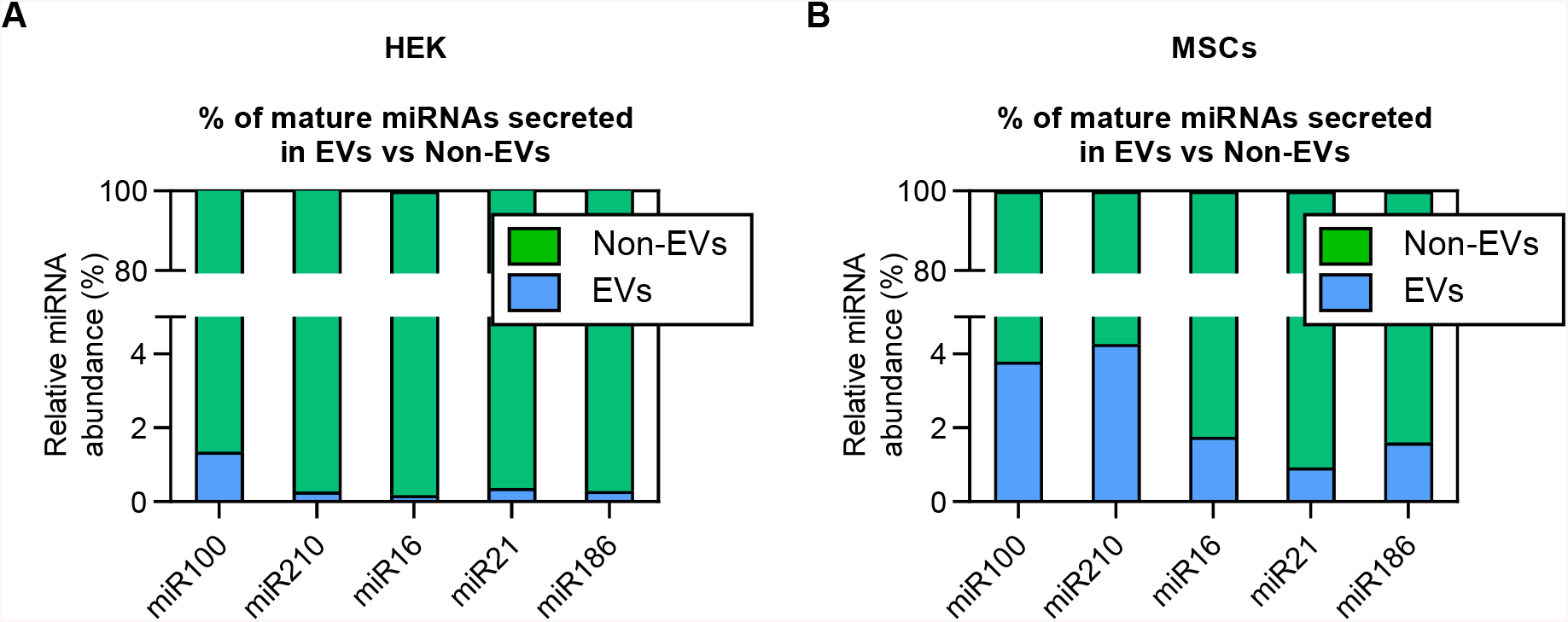
RT-qPCR quantification of mature miRNAs from EV and Non-EV fractions at their basal cellular expression level. Regardless of relative miRNA enrichment levels, the majority of secreted miRNAs are found outside of EVs both in HEK **(A)** and MSC **(B)** samples and the Non-EV material contains a striking 96–99% of total secreted miRNAs.

## DISCUSSION

The extracellular space consists of multiple types of nucleic acid carriers including EVs, which are one of the most actively studied fractions of the secretome. EVs contain many different types of RNA and as widely demonstrated, transfer it in a biologically active form^6,26,27^. Owing to the RNA transfer capacity, EVs are actively developed as carriers of biotherapeutic RNA molecules, such as miRNAs and shRNAs. Since the overall abundance of specific miRNA molecules in EVs is low and only a small fraction of all EVs contain a given miRNA sequence^12–14^, many efforts are directed towards enriching EVs with therapeutic miRNA and shRNA sequences, often by overexpressing them in EV producer cells. Owing to studies often using ultracentrifugation for EV isolation, it is known that certain RNA stretches are rather secreted independently of EVs^28,29^. Yet, it must be noted that incomplete EV pelleting in centrifugation-based approaches perplexes the interpretation of those results, indicating to the need of employing more refined methodologies for dissecting the secretome^30,31^.

The study at hand employed a well-established secretome fractionation strategy, SEC in order to explore the relative distribution of short RNA stretches in EVs and non-EV secretory fractions. Our preliminary observations in the RNA overexpression context showed that <2% of analysed miRNAs were secreted via EVs, while the great majority of overexpressed miRNA ended up in the non-EV fraction, corroborating with a recent study showing similar findings from virus infected cells^15^. Moreover, the low EV-miRNA amount was observed irrespective of the overexpression type, using either CMV or U6 promoter and pri-miRNAs/shRNAs for loading the EVs, as often used for therapeutic RNA loading into EVs^32–36^. This indicates that passive overexpression strategy could be a rather wasteful way for loading EVs with RNA cargo molecules. Also, though the overexpressed EV miRNA was fully processed and stable in serum-containing environment up to 24 h thereby greatly exceeding the half-life of EVs in vivo^5,37–39^, the low miRNA level is still unlikely to have satisfactory therapeutic potential.

Owing to the insufficient systematic knowledge on how short RNA sequences are secreted in the global context on the basal level, it is difficult to design robust universal strategies for miRNA enrichment into EVs. Though a number of small RNA sequencing studies have been performed to define the exact nucleic acid composition of EVs, the majority thus far exploit UC based EV enrichment protocols^14,40–42^. Further complexity in studying EV-RNA arises from the low overall abundance of specific miRNA molecules in EVs^12–15^ and the knowledge that sRNAs cover only the minority of reads, even when concentrating on studying short reads^23,24^. Another aspect in such global analysis is the presence of bovine serum-derived miRNAs, which can contaminate both EV and non-EV samples when EV-depleted serum is used^19,20^. Though this can successfully be overcome by using Opti-MEM based serum free media, it must be considered that altering the culturing conditions might change the transcriptome of cells and therefore also affect the contents of EVs, as seen previously^21^. To minimise the potential risks of changes in EV content and contaminating RNA originating from EV-depleted serum^19,20^, the study at hand conducted parallel comparisons (i.e. the cell, EV and non-EV samples were collected from within the same experiment) and used Opti-MEM based serum free media for EV production, respectively.

Hence, to follow up on the aforementioned overexpression work, we continued to use SEC, separated the secretome of HEK cells and MSCs into the EV and non-EV fractions and aimed to characterize the global sRNA sorting at the basal expression level. The preliminary clustering analysis showed a clear separation between cellular, EV and non-EV samples. While this suggested of global differences, the content of EVs and non-EVs appeared to be more similar to each other than to their source cells. A closer look at different short RNA biotypes revealed that the similarity might be attributed to piRNA sequences, of which the cells were relatively depleted of, as well as to miRNAs, being highly prevalent in cells and less abundant in the secretory fractions. Nevertheless, the miRNA expression both in EVs and non-EVs correlated highly to that of their parent cells, suggesting that in general the most important factor determining the level of secreted miRNA, is its expression level in the source cells. This does not however mean that miRNA sequences cannot be enriched in EVs, as certain sequence-specific miRNA sorting into EVs mediated by specific miRNA binding proteins such as hnRNP-Q, hnRNPA2B1 has been validated^43,44^. Similarly, to those observations, compared to source cells, we also noticed relative enrichment of certain miRNA sequences in EVs as well as in the non-EV fraction. Interestingly, these sequences were expressed in HEK cells and MSCs at a low level, whereas the secretion of highly-expressed miRNAs was generally not enriched.

Importantly, we must underline that even if a specific sequence was enriched in EVs vs non-EVs in relative terms, as the total abundance of miRNAs in the non-EV fraction in fact greatly exceeded that of EVs. Though similar evidence has been reported previously by others, often by using UC for EV isolation^28,29^, we were now able to confirm these observations on SEC-purified secretome fractions by using quantitative RT-qPCR. In the cell culture supernatants of HEK cells, we found less than 2% of miRNAs in EVs, whereas over 98% of miRNA was found in the non-EV fraction. MSC-derived samples contained slightly more miRNAs in EVs, however, over 96% of miRNAs were still secreted in non-EVs fraction. This could on one hand reflect the inherent differences in cell line dependent packaging mechanisms, else hint to mechanisms modulating the preservation of basal therapeutic potential of MSC EV miRNAs^45,46^. Moreover, we saw that the basal expression and secretory enrichment level of miRNAs is irrelevant to their overall accumulation in EVs, as the secretion is nonetheless heavily inclined towards the non-EV pool. Considering our aforementioned observation that even miRNA/shRNA overexpression is unable to reroute the miRNAs more to the EVs, transient overexpression strategy seems to be uneconomic for loading EVs with RNA cargo. Though sequences with low basal expression level ought to give better results in terms of expression level increase, and even if significant enrichment is achieved, the bulk material will still not end up in EVs. This overall pinpoint that without active EV-targeted loading strategies, the level of a given miRNA in the vesicles will most probably not reach to a therapeutically relevant scale.

In summary, this study provides important information on the basal- and overexpressed miRNA sorting patterns, outlining that secretory sRNA sorting is heavily inclined towards non-vesicular extracellular pool. Owing to the therapeutic interest in EV-associated sRNAs, it is increasingly important to understand and avoid non-vesicular secretion mechanisms. While numerous steps to tackle obstacles of therapeutic sRNA loading into EVs have been made, need for future strategies that finely alter the secretion pattern of short RNA molecules still remains.

## METHODS

### miRNA overexpression

To express precursors for the specific miRNAs, pri-miRNA sequences were amplified by PCR using Phusion® High-Fidelity DNA Polymerase (NEB BioLabs, USA) from human embryonic kidney (HEK293T) cells genomic DNA with the oligonucleotide primers described by Diederichs *et al* ^47^ and purchased from Integrated DNA Technologies (USA). The obtained pri-miRNA sequences were cloned into the pEGFP-N1 backbone vector by using AgeI and XbaI restriction enzymes and T4 DNA ligase (NEB BioLabs, USA). For shRNA overexpression, luciferase-targeting shRNA oligonucleotides (Integrated DNA Technologies, Belgium) were cloned promotor into the shRNA expression plasmid, downstream of the U6 promoter, using the BbsI restriction enzyme^32^. The shRNA oligonucleotide sequences were: CCGGAAGAGCACTCTCATCGACAAGCTCGAGCTTGTCGATGAGAGTGCTCTTTTTTTG, and AATTCAAAAAAAGAGCACTCTCATCGACAAGCTCGAGCTTGTCGATGAGAGTGCTCTT Subcloning Efficiency™ DH5α™ Competent Cells (Life Technologies Invitrogen, USA) were used to propagate the plasmid, followed by plasmid purification using QIAprep Spin Miniprep Kit (Qiagen, Germany) and sequencing (Source BioScience – Life Sciences, UK). For overexpression studies HEK293T cells were transfected with pri-miRNA expression vectors by using polyethyleneimine (PEI, Cat.No. 408727, Sigma-Aldrich) transfection at a DNA:PEI ratio of 1:4. After 4 h post-transfection, the cells were washed once with phosphate buffered saline (PBS) solution (Thermo Fisher Scientific, Waltham, MA, USA) followed by media changed to OptiMEM Reduced serum medium (Thermo Fisher Scientific, Waltham, MA, USA). 48h postconditioning, the EVs were isolated as described below.

### EV isolation for overexpression and validation

For miRNA overexpression studies, after 48h of media conditioning, the EVs were isolated by using the Amicon Ultra-15 100 kDa molecular weight cut-off spin-filters (Millipore, Germany) discriminating the retentate and flow-through/filtrate. Else, the retentate was subjected to ultrafiltration SEC fractionation using the S-400 as described in^18^(fractionation of overexpressed miRNA secretome) or fractionated by Tricorn 10/300 columns (Sigma-Aldrich, Saint-Louis, MO, USA) packed in-house with Sepharose Fast Flow 4 resin (GE Healthcare, Pittsburgh, PA, USA) (miRNA validation). In case of miRNA overexpression, the A280 and RNA quantification measurements (Quant-iT RiboGreenRNA Assay Kit; Thermo Fisher Scientific, Waltham, MA, USA) across the elution profile were used to set the location of fractions 1 to 5 (each spanning 10ml) in a way that the cross-talk between the set ranges would be minimal, yet would cover the major changes in the protein and RNA profiles. For miRNA validation, the ‘non-EV’ fraction contained the non-vesicular fractions from the SEC as well as the flow-through from the 100kDa ultrafiltration step.

### miRNA quantification by qRT-PCR

For miRNA quantification, an appropriate amount of Trizol LS (Thermo Fisher Scientific, Waltham, MA, USA) was used to prepare the samples for subsequent RNA extraction according to the manufacturers’ protocol. For miRNA validations, the variation in RNA extraction efficiencies, 3 μl of a 5 μM synthetic miRNA oligonucleotide, cel-miR-39 (5′-UCACCGGGUGUAAAUCAGCUUG) (IDT, Leuven, Belgium), was added to each sample at the phenol extraction stage. In all cases, cDNA synthesis was performed by using the TaqMan MicroRNA Reverse Transcription Kit and the respective hairpin primers from TaqMan MicroRNA Assays (let-7a-5p assay ID:000377 and let-7b-5p assay ID:000378, miR-186-5p assay ID:002285; miR-210-3p assay ID: 000512; miR-16-5p assay ID:000391; miR-21-5p assay ID:000397 and miR-100-5p assay ID:000437; all from Thermo Fisher Scientific, Waltham, MA, USA). The quantitation was performed by using the TaqMan Gene Expression Master Mix together with the respective TaqMan MicroRNA Assays on a Step-One Real-Time PCR instrument (Thermo Fisher Scientific, Waltham, MA, USA). shRNA detection was performed by synthesising cDNA as above, but using custom-designed hairpin RT primers, and amplified using KiCqStart SYBR Green qPCR Ready Mix (Sigma Aldrich) on a Step-One Real-Time PCR instrument as above. The PCR efficiency of each reaction was calculated with LinRegPCR program (developed by Dr Jan Ruijter, Heart Failure Research Center, The Netherlands), while the Ct values were obtained from StepOne Software (Applied Biosystems, USA). In the case of overexpression studies, an equal amount of input RNA was used for reverse transcription. For miRNA validations, the total miRNA quantity was normalized to cel-miR-39 levels and further corrected for the volume of the respective fractions (Sepharose Fast Flow 4 fractionation volumes of 3-3.5 ml for EVs and 131-133 ml for non-EV (non-EV fraction + flow through of 100kDa ultrafiltration units)). For EV miRNA stability studies, EVs were isolated and purified as above, mixed with FBS at a final serum concentration of 10% and incubated at 37°C for indicated times. At the end of each time point, Trizol LS was added to samples and the RNA was extracted and analysed by qPCR as above.

### EV isolation for small RNA sequencing

Neuro2a, MSC and HEK cell-lines were maintained in high-glucose (4.5 g/l) DMEM GlutaMax (Thermo Fisher Scientific, Waltham, MA, USA; HEK) or RPMI medium 1640 (Thermo Fisher Scientific, Waltham, MA, USA; MSCs) supplemented with 10% fetal bovine serum (FBS, Thermo Fisher Scientific, Waltham, MA, USA). All cells were cultured in a humidified atmosphere at 37°C and 5% CO_2_. 6×10^6^ cells were first seeded onto 15 cm cell culture dishes. After 24 hours, cells were washed with 0.01M phosphate buffered saline (PBS) to ensure removal of any residual FBS before a media change to OptiMEM (Thermo Fisher Scientific, Waltham, MA, USA) supplemented with 1% Antibiotic Antimycotic Solution (Sigma-Aldrich, Saint-Louis, MO, USA). The supernatant was collected after 48 hours and EVs were harvested by ultrafiltration size exclusion liquid chromatography fractionation as described in^18^. To obtain the non-vesicular pool, flow-through from the 100kDa ultrafiltration step was combined with the non-vesicular fractions from the SEC fractionation and thereafter concentrated on the 10-kDa molecular weight cut-off (MWCO) Amicon spin filters (Merck Millipore), the latter of which was also used to concentrate the vesicular fractions. In some control experiments, when specifically indicated, 100 kDa MWCO spin filters were used instead. Immediately after concentrating, all samples were mixed with Trizol LS (Thermo Fisher Scientific, Waltham, MA, USA) and stored at −80°C until RNA extraction for small RNA sequencing analysis. The abovementioned workflow was used to generate three biological replicates from each cell line.

### EV characterization

Nanoparticle tracking analysis (NTA) was performed by using the NanoSight NS500 nanoparticle analyser (Malvern Instruments, Malvern, UK) to measure the size distribution and concentration of the purified EVs. The EV samples were diluted in PBS and the movement of the particles was recorded within five 30 seconds videos that were subsequently analysed using the NTA Software 2.3 (Malvern Instruments, Malvern, UK) at detection threshold 5, camera gain 350 and shutter setting 800. Transmission Electron Microscopy (TEM) was performed as described in^48^.

### Sample preparation for small RNA sequencing

Details of sample preparation can be found in ^23^. Briefly 60 ng ±10% of total RNA was subjected to small RNA library preparation by using the NEBNext Multiplex Small RNA Library Prep Set 1 and 2 for Illumina (NEB, Ipswich, MA, USA) kit according to the manufacturer’s instructions. After quantification with the KAPA Library Quantification Kit (Cat.No. KK4824, Kapa Biosystems, London, UK), samples were pooled at the equimolar ratio. Two libraries were generated in parallel, each one eventually containing a pool of 18 barcoded samples. The readymade libraries were further checked on the High Sensitivity D1000 ScreenTape (Agilent Technologies, Lanarkshire, UK) and quantified by using the KAPA Library Quantification Kit to enable precise loading of the flow cell. The clusters were generated by using the cBot and two flow cells (4 lanes; 2 technical replicates per library) were sequenced on the HiSeq2500 (HiSeq Control Software 2.2.58/RTA 1.18.64, Illumina Inc., San Diego, CA, USA) with a 1×51 setup in RapidRun mode using ‘HiSeq Rapid SBS Kit v2’ chemistry.

### Small RNA sequencing data analysis

Small RNA sequencing data analysis regarding mapping and annotation was performed as outlined in^23^. For data visualization (principle component analysis, correlation scatterplots, heatmaps), raw read counts were transformed in R-studio software (R version 3.4.2)^49^ by using the approach of variance stabilizing transformation (vsd; blinded=F). Differential expression analysis of miRNAs was performed by using the R package DESeq2^50^, including mature miRNAs exceeding a raw read count cut-off across the studied samples (RowMeans >1 within the studied cell line). Heatmaps of differentially expressed miRNAs (vsd counts, local zero-centred normalization) were generated by using the Multiple Experiment Viewer program (Version 4.9.0)^51^. The miRNAs were hierarchically clustered by using the average linkage clustering method, Pearson distance metric and optimized gene leaf order.

### Statistical analysis

Statistical fold enrichment analyses were performed using GraphPad Prism Version 7 (GraphPad Software, Inc., La Jolla, CA, USA) by using the two-way Anova with Tukey post-hoc correction. The statistics from the differential expression analysis of miRNAs were generated by the R software package DESeq2^50^.

## ACKNOWLEDGEMENTS

The authors would like to acknowledge support from Science for Life Laboratory, the National Genomics Infrastructure, NGI and Uppmax for providing assistance in massive parallel sequencing and computational infrastructure. We would also like to thank Assistant Professor Marc Friedländer and Wenjing Kang for their assistance in the bioinformatic data analysis.

## AUTHOR CONTRIBUTIONS

I.M. and S.E.A. conceived the idea of the study; H.S., K.K., I.M. and S.E.A. designed the experiments; I.M., G.C., J.Z.N, Y.X.F.L and H.S. isolated the EVs for sequencing; M.C., G.C., I.M., P.V., M.P. and N.P performed EV characterisation, qRT-PCR and other validation experiments; H.S. and K.K. prepared the sequencing libraries; H.S. performed and J.O.W. advised with bioinformatic analysis; G.C. acquired TEM images; S.E.A. and M.J.W. gave guidance throughout the study; H.S. and I.M. analysed and compiled the data, and wrote the manuscript.

