## Supplementary data for "Profiling of extracellular small RNAs highlights a strong bias towards non-vesicular secretion"

[#] Shared authorship

* To whom correspondence should be addressed. Tel: +46858583657; Fax: +46 8 585 838 00; (H.S.) and Tel: 01865272199; Fax: 01865272420; (I.M.)

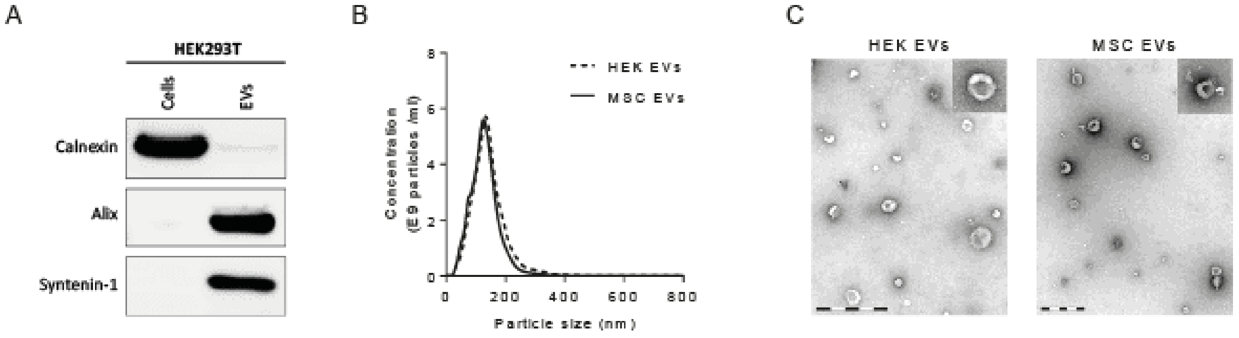


**Supplementary Figure 1. EV characterization** (A) A representative Western blot demonstrating the presence of EV marker proteins Alix and Syntenin-1 in isolated EV samples. (B) Representative nanoparticle tracking analysis profiles demonstrating the size distribution profile of isolated EV samples. (C) Representative electron microscopy images demonstrating the typical EV size and morphology in isolated samples. Scale bar 500nm.

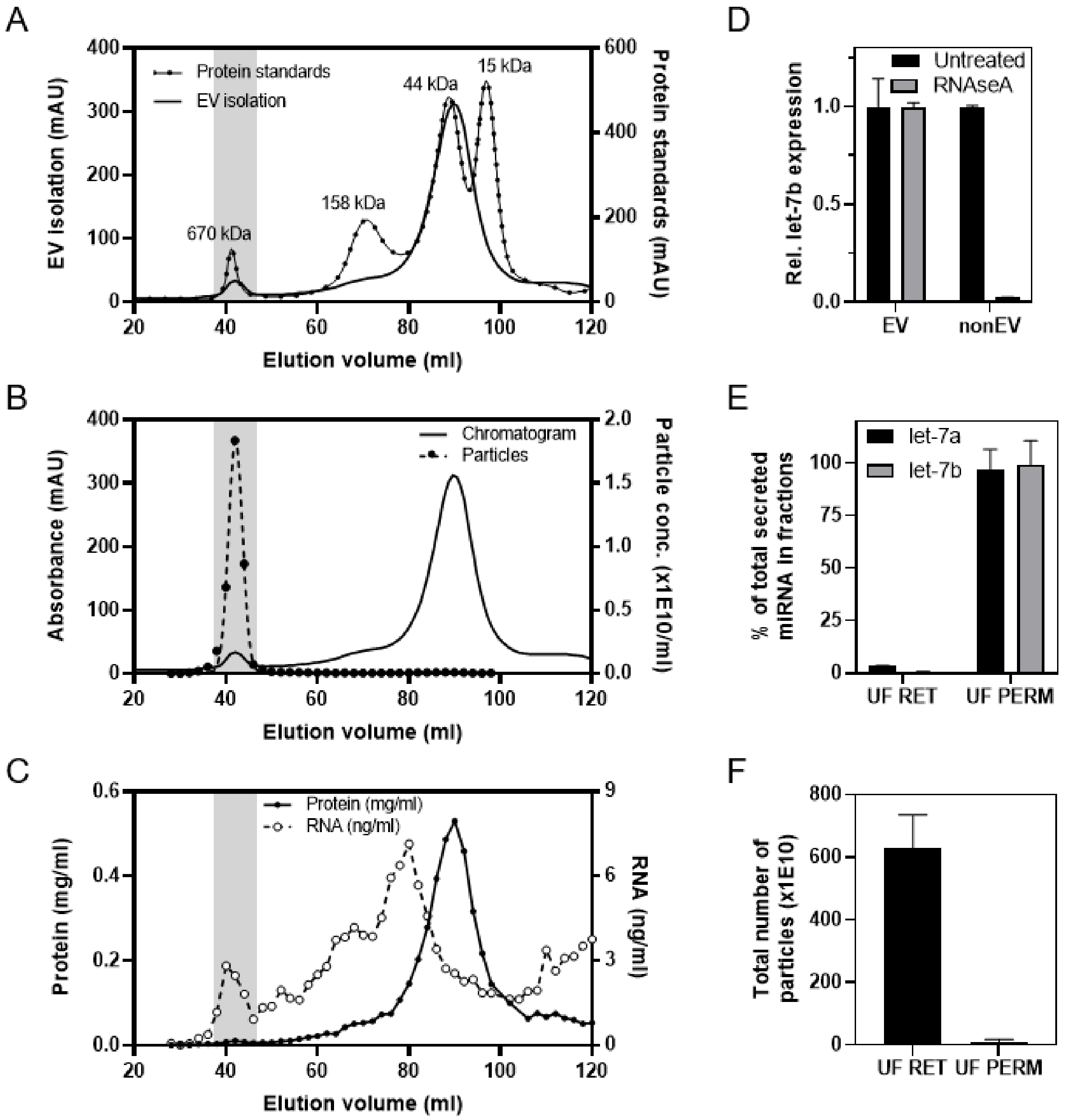


**Supplementary Figure 2. Further validation of the SEC workflow for separating EV and non-EV secreted material using Neuro2a cells.** The SEC chromatograms (A, B, C) indicate that the EV peak (i.e. the void volume peak) contains material > 600 kDa MW and is the only fraction to contain particles, as detected by the Nanoparticle Tracking Analysis. The elution profile of RNA and protein (C) is more complex, where a significant proportion of material is found in the non-EV area of the chromatogram. (D) miRNA in the EV fraction is protected from RNAseA treatment, whereas the non-EV fraction was susceptible for that. (E) When the conditioned medium is concentrated with 100 kDa MWCO spin filters the majority of miRNA is found in the ultrafiltration permeate (UF PERM), which is consistent with the observation that the majority of miRNAs are found in the non-EV peak of the SEC workflow. Ultrafiltration retentate (UF RET) contained only a minor fraction of miRNA). (F) No particles were detected in the UF permeate, again in line with the observation that the majority of miRNA is found in the non-EV fraction.

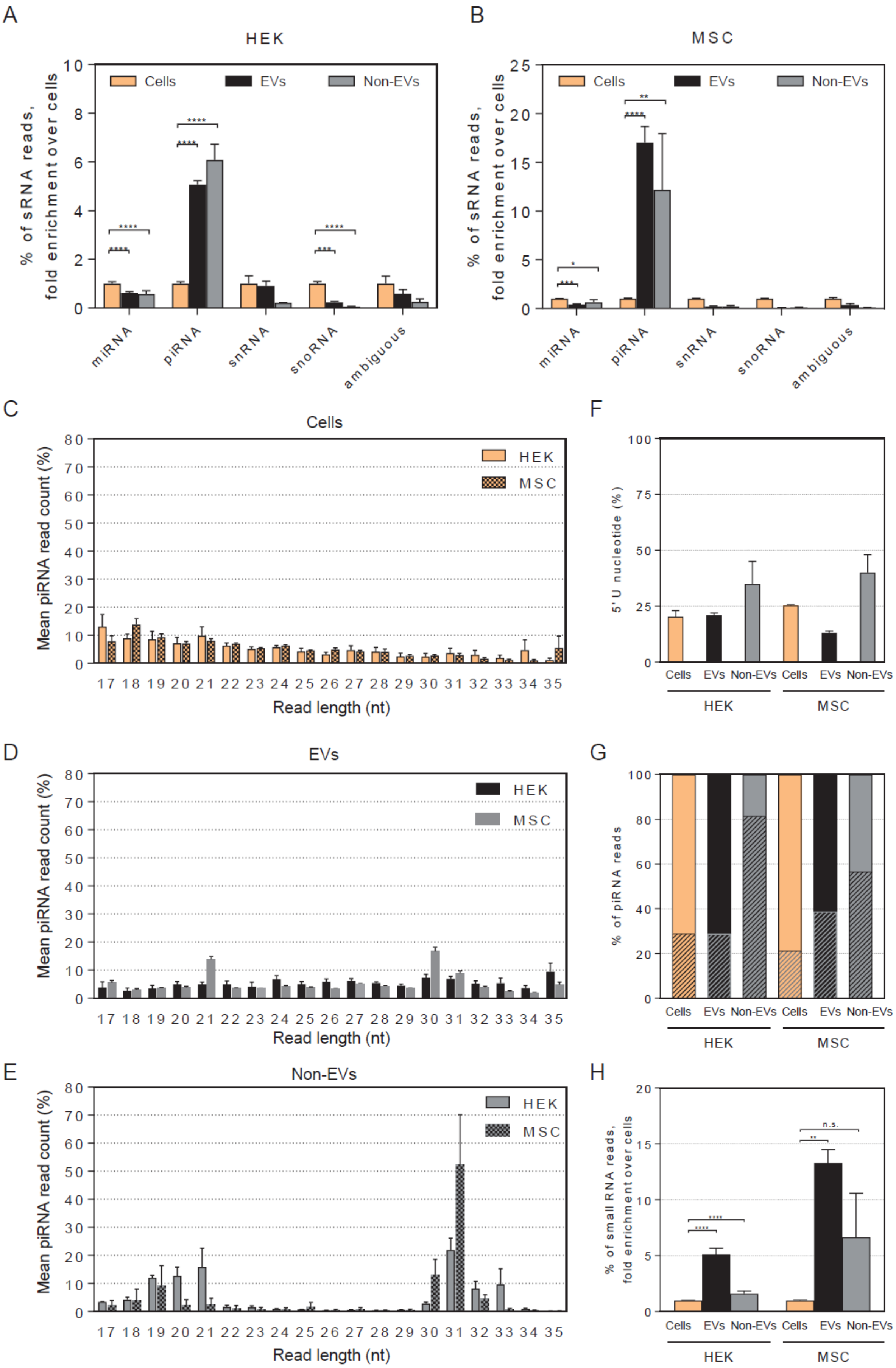
A
HEK
% of sRNA reads,
fold enrichment over cells
Cells
EVs
Non-EVs
miRNA
piRNA
snRNA
snoRNA
ambiguous
B
MSC
C
Cells
Mean piRNA read count (%)
HEK
MSC
Read length (nt)
D
EVs
E
Non-EVs
F
5' U nucleotide (%)
G
% of piRNA reads
H
% of small RNA reads,
fold enrichment over cells
n.s.

**Supplementary Figure 3. sRNA composition, characteristics and enrichment in secretory fractions.** The percentage of sRNA biotype reads, fold enrichment over cells in HEK (A) and MSC (B) samples reveals that EV and Non-EV samples are enriched in piRNA sequences and depleted from miRNA sequences. piRNA reads distribution length in cells (C), EVs (D) and Non-EVs (E), and the presence of 5' U (F) suggests that a proportion of genes annotated as piRNAs might not be true piRNAs. Up to a third of piRNA annotations in cells and EVs, and up to 80% of piRNA annotations in Non-EVs also align to tRNA reads (G). Even if the tRNA-overlapping piRNA sequences are eliminated from the analysis, the enrichment of piRNA sequences in EVs and Non-EVs is still observed.

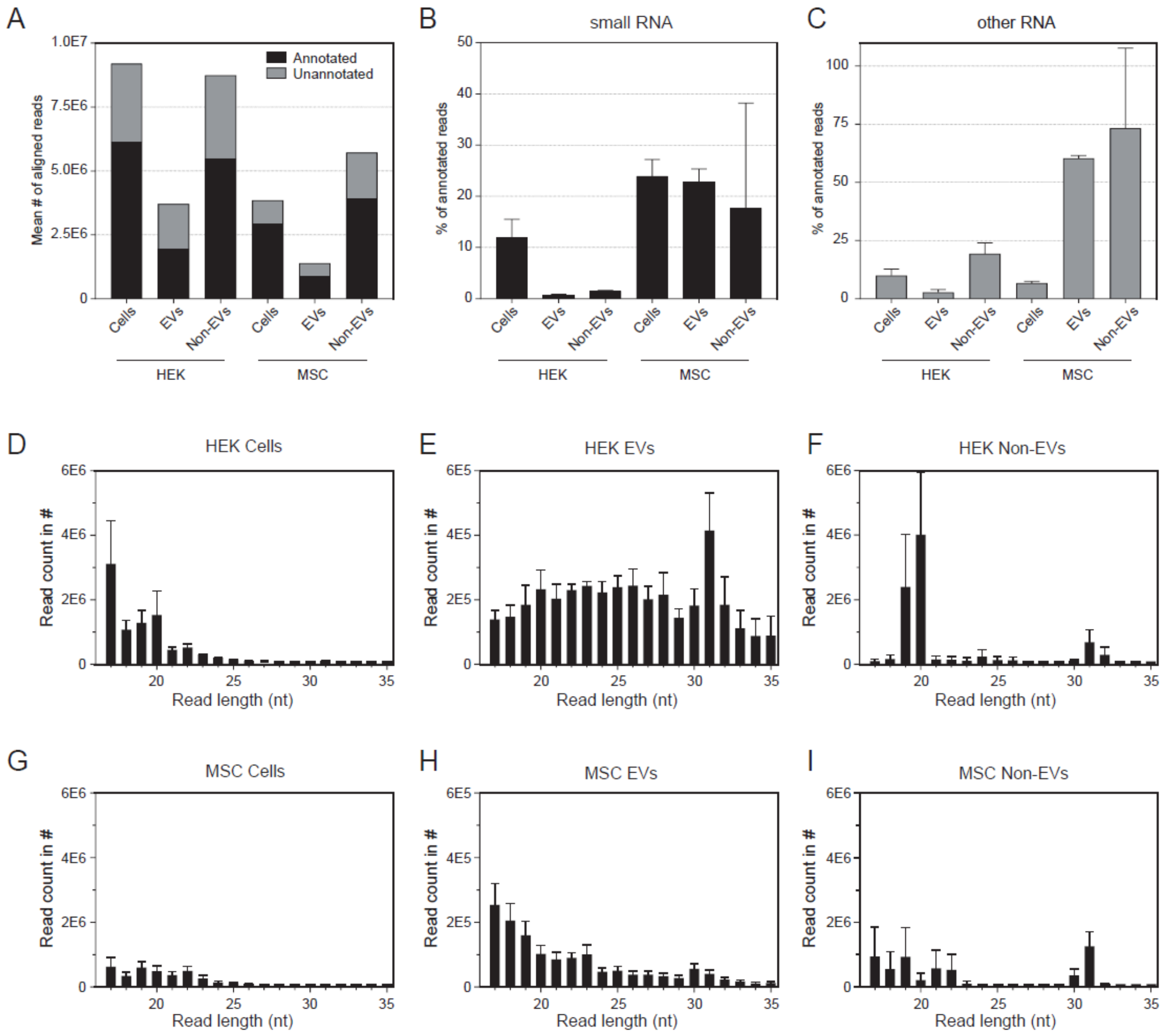


**Supplementary Figure 4. Read alignment, annotation and length statistics across the studies samples.** The average number of aligned reads varies between sample types (A). The percentage of annotated reads that align to sRNA biotype RNAs and other RNA sequences is presented in panels (B) and (C), respectively (respective rRNA data is not presented). Length distribution histograms of all annotated reads (sRNA, rRNA and other RNA) for HEK and MSC cells, EVs and Non-EVs is presented in panels (D–I).

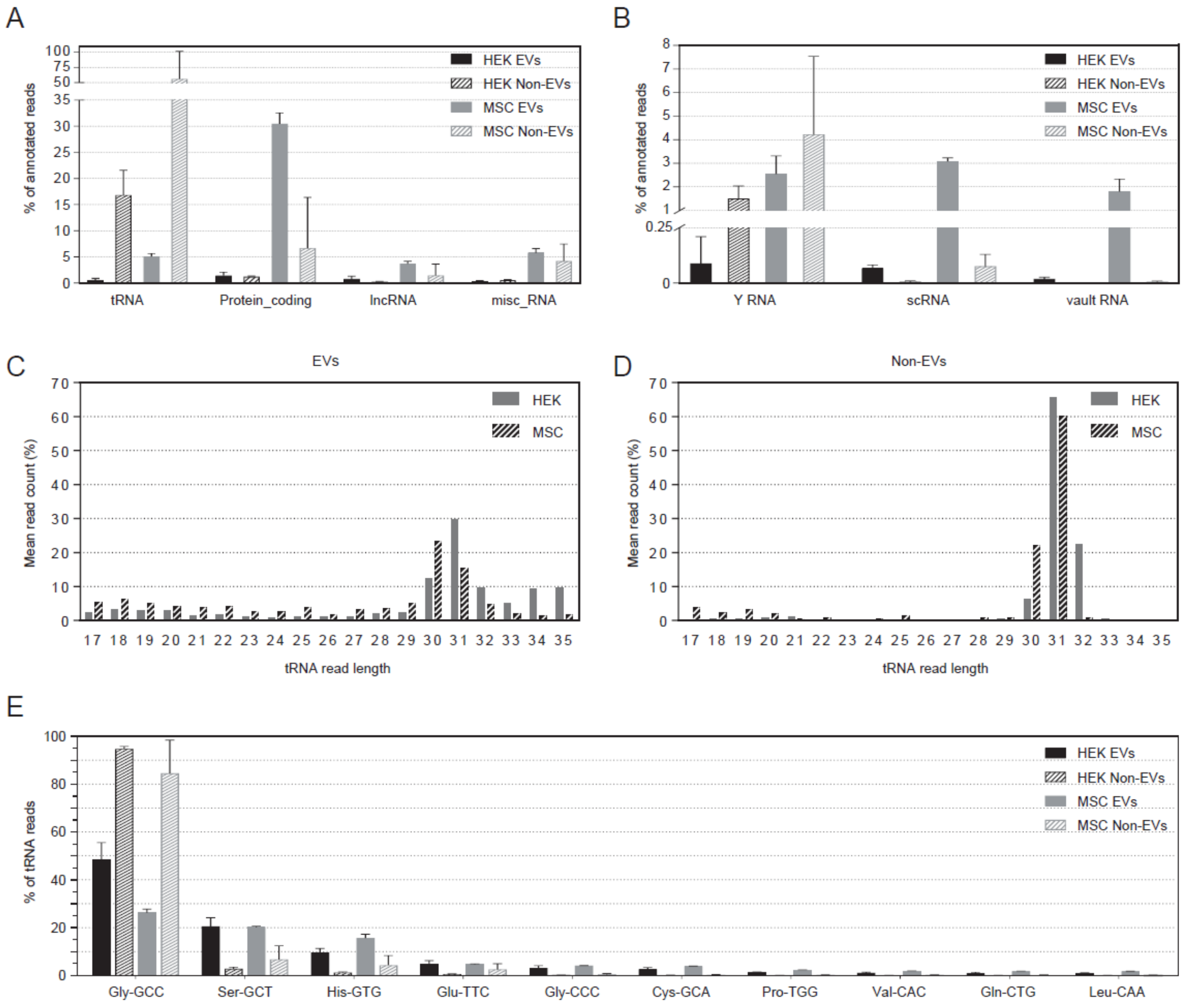


**Supplementary Figure 5. The prevalence of 'other RNA' biotypes** (A) The percentage of annotated reads aligned to tRNA, protein coding RNA, long non-coding RNA (lncRNA) and miscellaneous RNA (misc_RNA). (B) The percentage of annotated reads aligned to Y RNA, small cytoplasmic RNA (scRNA) and vault RNA. (C, D) Read length distribution histograms of tRNA-aligned reads in HEK and MSC EVs and Non-EVs. (E) Among the tRNA-aligning reads, Gly-GCC tRNA sequences are overrepresented, constituting up to 90% of all detected RNA reads in the Non-EV samples.

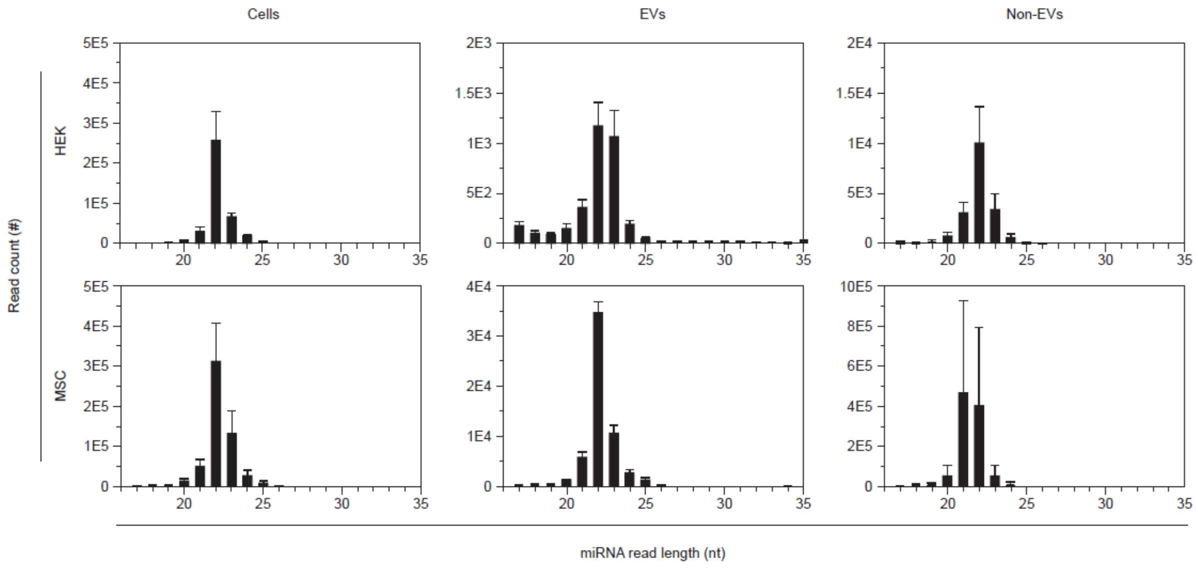


**Supplementary Figure 6. Length distribution histograms for reads aligning to miRNAs.** Sequences in all samples are in accordance with an expected length of mature miRNAs.

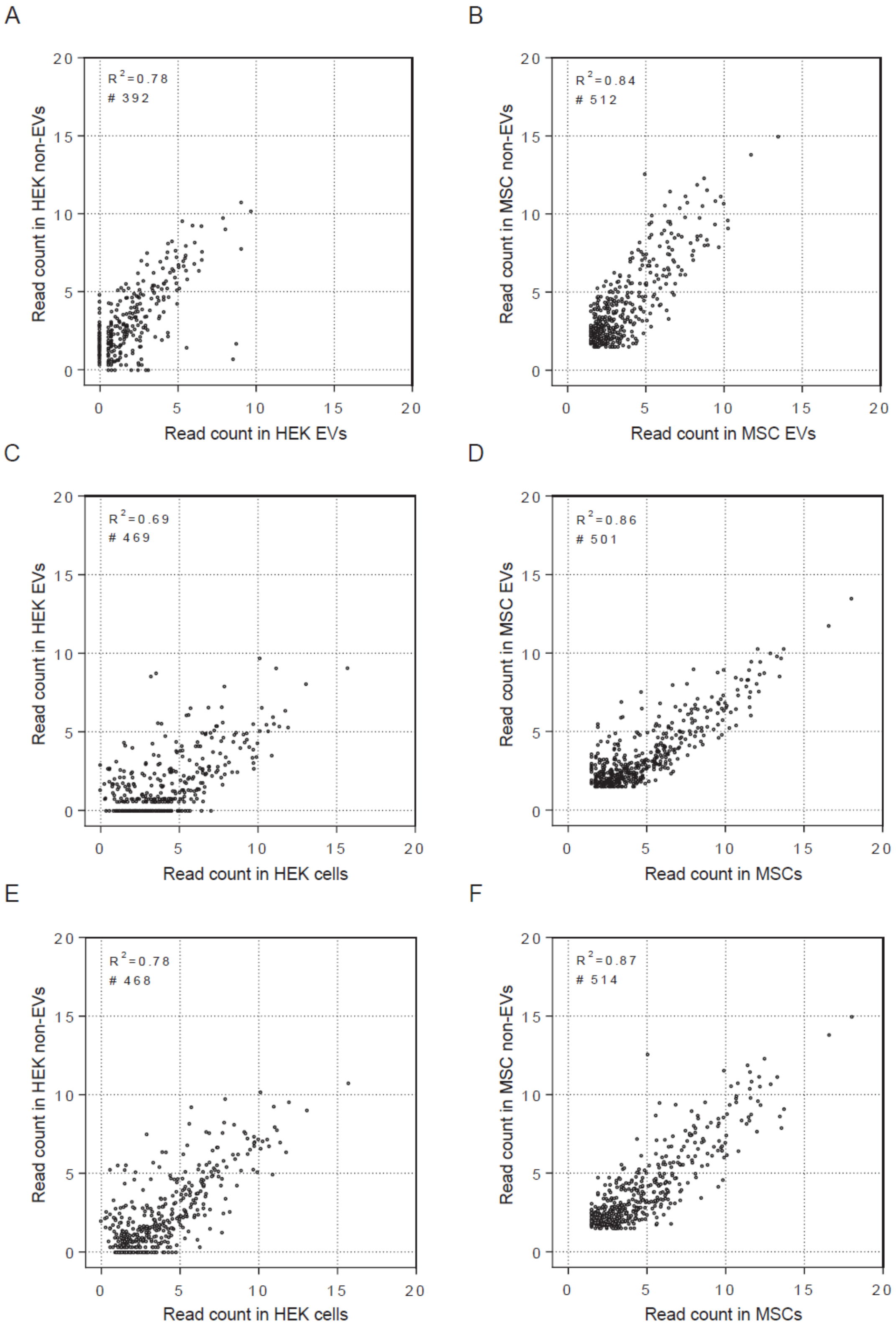


**Supplementary Figure 7. Correlation plots of all mature miRNAs.** miRNA expression profile between EV, Non-EV and cell samples are highly correlated both in HEK (A, C, E) and MSC (B, D, F) samples.

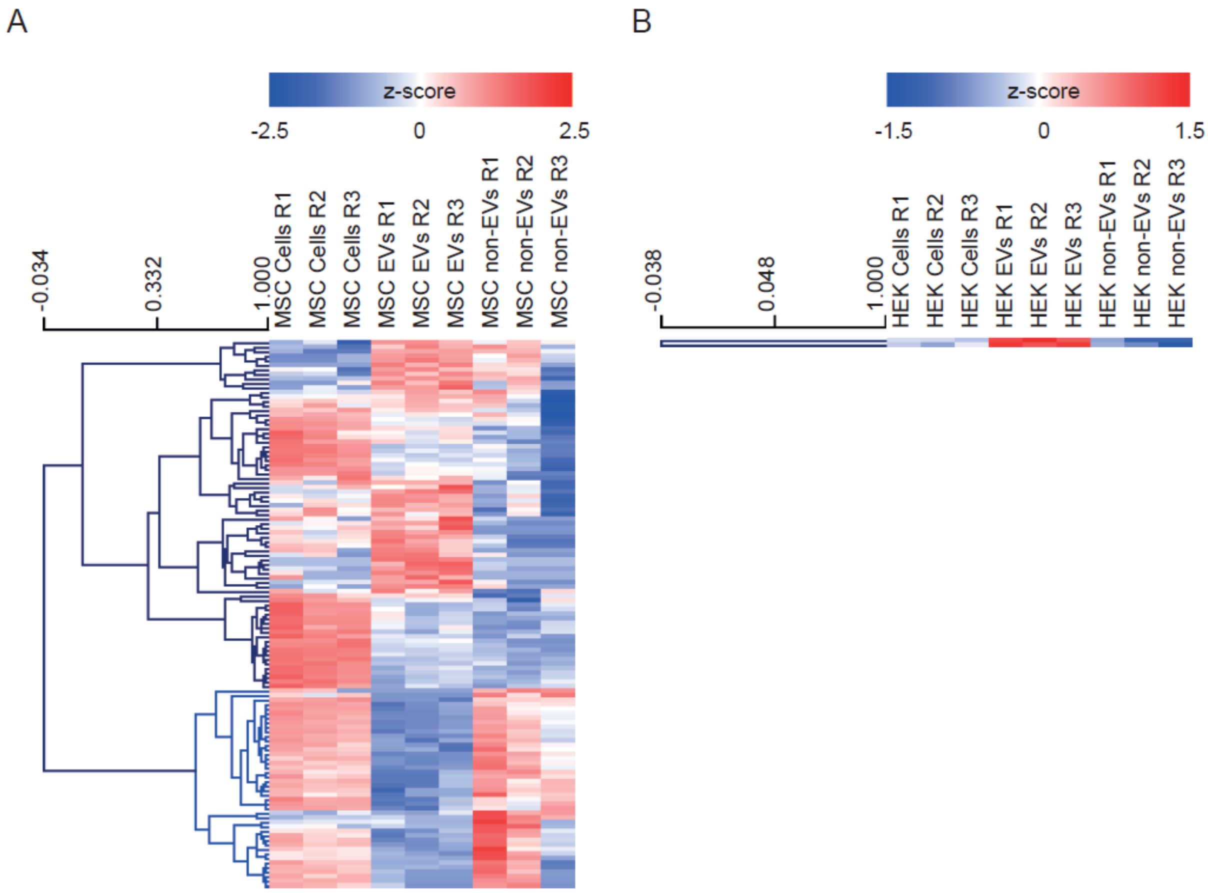


**Supplementary Figure 8. Heatmaps of differentially expressed miRNAs detected between the EV and Non-EV fraction of the MSC (A) and HEK (B) secretome.** EV-enriched miRNAs tend to have low expression level in cells, whereas EV-depleted miRNAs are sampled from miRNAs with high cellular expression. In HEK samples, there are only two miRNAs that are differentially expressed between EV and Non-EV samples (B), whereas in MSC samples the number of differentially expressed miRNAs is considerably higher.

**Supplementary Table 1. Differentially expressed miRNAs in EVs *vs* Cells.**

| EV *versus* cells | | | | | | | | | |
|---|---|---|---|---|---|---|---|---|---|
| DE miRNA name | HEK | | | | DE miRNA name | MSC | | | |
| | log2FC | lfcSE | pvalue | padj | | log2FC | lfcSE | pvalue | padj |
| hsa-miR-7704 | 7.9 | 1.0 | 3.34E-14 | 9.97E-12 | hsa-miR-6892-5p | 7.1 | 0.7 | 4.5E-16 | 3.4E-14 |
| hsa-miR-6087 | 7.3 | 1.0 | 1.21E-12 | 1.80E-10 | hsa-miR-365a-5p | 6.4 | 0.7 | 2.0E-18 | 3.8E-16 |
| hsa-miR-4492 | 6.0 | 1.2 | 1.74E-06 | 6.47E-05 | hsa-miR-128-1-5p | 6.2 | 0.9 | 4.1E-10 | 1.5E-08 |
| hsa-miR-4497 | 5.9 | 1.2 | 4.87E-07 | 3.63E-05 | hsa-miR-503-3p | 6.2 | 1.0 | 1.4E-07 | 2.6E-06 |
| hsa-miR-3610 | 5.8 | 1.1 | 1.26E-06 | 6.28E-05 | hsa-miR-193b-5p | 5.7 | 0.7 | 1.4E-17 | 1.4E-15 |
| hsa-miR-4508 | 5.5 | 1.2 | 1.64E-06 | 6.47E-05 | hsa-miR-451a | 5.7 | 0.9 | 3.1E-09 | 9.1E-08 |
| hsa-miR-4792 | 5.4 | 1.1 | 1.40E-08 | 1.39E-06 | hsa-miR-365b-5p | 5.7 | 0.6 | 9.9E-20 | 3.8E-17 |
| hsa-miR-1225-3p | 4.9 | 1.3 | 1.45E-04 | 3.09E-03 | hsa-miR-6807-5p | 5.6 | 1.0 | 2.5E-07 | 4.4E-06 |
| hsa-miR-149-3p | 4.5 | 1.3 | 3.85E-04 | 6.75E-03 | hsa-miR-1307-5p | 5.5 | 0.6 | 3.9E-18 | 5.1E-16 |
| hsa-miR-199a-3p | 4.3 | 0.9 | 1.15E-06 | 6.28E-05 | hsa-miR-323a-5p | 5.4 | 0.9 | 2.4E-08 | 5.5E-07 |
| hsa-miR-3648 | 4.2 | 1.1 | 2.27E-06 | 7.52E-05 | hsa-miR-152-5p | 5.3 | 0.7 | 1.2E-11 | 5.3E-10 |
| hsa-miR-654-5p | 4.0 | 1.3 | 8.13E-05 | 2.02E-03 | hsa-miR-1225-3p | 5.1 | 1.0 | 1.2E-05 | 1.4E-04 |
| hsa-miR-199b-3p | 3.7 | 0.9 | 3.49E-05 | 1.04E-03 | hsa-miR-410-5p | 4.9 | 1.0 | 1.5E-05 | 1.8E-04 |
| hsa-miR-4787-3p | 3.5 | 1.2 | 4.04E-03 | 3.65E-02 | hsa-miR-23a-5p | 4.9 | 0.8 | 8.6E-10 | 2.8E-08 |
| hsa-miR-4516 | 3.5 | 1.3 | 3.52E-04 | 6.55E-03 | hsa-miR-433-5p | 4.8 | 1.1 | 1.1E-05 | 1.4E-04 |
| hsa-miR-365b-5p | 3.5 | 1.2 | 3.66E-03 | 3.64E-02 | hsa-miR-7977 | 4.6 | 0.8 | 2.4E-08 | 5.5E-07 |
| hsa-miR-940 | 3.5 | 1.2 | 2.00E-03 | 2.29E-02 | hsa-miR-6892-3p | 4.6 | 1.2 | 2.7E-04 | 2.2E-03 |
| hsa-miR-143-5p | 3.4 | 1.3 | 4.19E-03 | 3.67E-02 | hsa-miR-142-5p | 4.5 | 0.9 | 1.3E-05 | 1.5E-04 |
| hsa-miR-3195 | 3.4 | 1.5 | 5.61E-03 | 4.40E-02 | hsa-miR-92a-1-5p | 4.5 | 0.8 | 2.9E-08 | 6.3E-07 |
| hsa-miR-146a-5p | 3.3 | 1.0 | 7.47E-04 | 1.11E-02 | hsa-miR-4669 | 4.5 | 1.0 | 1.6E-05 | 1.8E-04 |
| hsa-let-7b-5p | 3.3 | 0.9 | 8.08E-05 | 2.02E-03 | hsa-miR-370-5p | 4.4 | 0.8 | 1.2E-07 | 2.3E-06 |
| hsa-miR-214-3p | 3.2 | 1.2 | 2.94E-03 | 3.22E-02 | hsa-miR-6789-3p | 4.4 | 1.1 | 5.8E-04 | 4.0E-03 |
| hsa-miR-4532 | 3.2 | 1.2 | 1.45E-04 | 3.09E-03 | hsa-miR-1275 | 4.3 | 0.8 | 6.0E-07 | 9.6E-06 |
| hsa-miR-379-5p | 3.2 | 1.1 | 1.91E-03 | 2.27E-02 | hsa-miR-149-3p | 4.2 | 1.0 | 1.8E-04 | 1.5E-03 |
| hsa-miR-125b-1-3p | 3.1 | 1.1 | 3.02E-03 | 3.22E-02 | hsa-miR-1226-5p | 4.2 | 1.2 | 4.1E-04 | 3.0E-03 |
| hsa-miR-143-3p | 3.0 | 1.0 | 6.51E-04 | 1.02E-02 | hsa-miR-194-3p | 4.1 | 1.2 | 1.2E-03 | 7.9E-03 |
| hsa-miR-206 | 2.9 | 1.0 | 1.51E-03 | 1.87E-02 | hsa-miR-431-3p | 4.0 | 0.8 | 5.3E-07 | 8.8E-06 |
| hsa-miR-21-5p | 2.5 | 0.9 | 3.99E-03 | 3.65E-02 | hsa-miR-3187-3p | 3.9 | 0.9 | 4.4E-05 | 4.1E-04 |
| hsa-miR-100-5p | 2.5 | 0.9 | 3.96E-03 | 3.65E-02 | hsa-miR-329-5p | 3.8 | 1.1 | 2.1E-03 | 1.2E-02 |
| hsa-miR-199a-5p | 2.5 | 0.9 | 5.00E-03 | 4.03E-02 | hsa-miR-487b-5p | 3.8 | 1.0 | 7.9E-04 | 5.3E-03 |
| hsa-let-7i-5p | 2.4 | 0.9 | 4.96E-03 | 4.03E-02 | hsa-miR-5100 | 3.8 | 1.0 | 4.1E-04 | 3.0E-03 |
| hsa-miR-432-5p | 2.3 | 1.4 | 3.49E-03 | 3.58E-02 | hsa-miR-92b-5p | 3.8 | 0.6 | 9.1E-11 | 3.5E-09 |
| hsa-miR-128-3p | -2.9 | 0.9 | 1.40E-03 | 1.81E-02 | hsa-miR-193a-5p | 3.7 | 0.5 | 6.3E-12 | 3.5E-10 |
| hsa-miR-103a-3p | -2.9 | 0.9 | 8.69E-04 | 1.23E-02 | hsa-miR-23b-5p | 3.7 | 0.8 | 3.6E-05 | 3.5E-04 |
| hsa-miR-148a-3p | -3.5 | 1.0 | 4.25E-04 | 7.04E-03 | hsa-miR-4443 | 3.7 | 0.9 | 4.5E-05 | 4.2E-04 |
| hsa-miR-615-3p | -3.6 | 1.0 | 3.16E-04 | 6.28E-03 | hsa-miR-486-5p | 3.6 | 0.8 | 2.4E-06 | 3.5E-05 |
| hsa-miR-629-5p | -3.7 | 1.3 | 4.40E-03 | 3.75E-02 | hsa-miR-4523 | 3.5 | 1.2 | 3.0E-03 | 1.6E-02 |
| hsa-miR-192-5p | -4.1 | 1.2 | 9.12E-04 | 1.24E-02 | hsa-miR-1343-3p | 3.4 | 0.9 | 2.0E-04 | 1.6E-03 |
| | | | | | hsa-miR-148b-5p | 3.4 | 0.9 | 4.4E-04 | 3.1E-03 |
| | | | | | hsa-miR-30b-3p | 3.3 | 1.1 | 5.2E-03 | 2.7E-02 |
| | | | | | hsa-miR-150-5p | 3.3 | 1.2 | 2.3E-03 | 1.3E-02 |
| | | | | | hsa-miR-4426 | 3.3 | 1.3 | 9.2E-03 | 4.4E-02 |
| | | | | | hsa-miR-654-5p | 3.2 | 0.5 | 1.2E-11 | 5.3E-10 |
| | | | | | hsa-miR-25-5p | 3.2 | 0.7 | 7.6E-06 | 1.0E-04 |
| | | | | | hsa-miR-125a-3p | 3.2 | 0.6 | 9.0E-08 | 1.8E-06 |
| | | | | | hsa-miR-877-5p | 3.1 | 1.3 | 8.3E-03 | 4.0E-02 |
| | | | | | hsa-miR-3187-5p | 3.1 | 1.3 | 1.0E-02 | 4.8E-02 |
| | | | | | hsa-miR-29b-1-5p | 3.0 | 1.0 | 3.7E-03 | 1.9E-02 |
| | | | | | hsa-miR-485-5p | 3.0 | 0.6 | 1.6E-06 | 2.5E-05 |
| | | | | | hsa-miR-1468-5p | 2.9 | 0.9 | 1.8E-03 | 1.1E-02 |
| | | | | | hsa-miR-1908-5p | 2.8 | 0.7 | 3.4E-05 | 3.3E-04 |
| | | | | | hsa-miR-541-3p | 2.8 | 1.0 | 7.4E-03 | 3.7E-02 |
| | | | | | hsa-miR-1228-5p | 2.6 | 1.1 | 9.3E-03 | 4.4E-02 |
| | | | | | hsa-miR-377-5p | 2.5 | 0.8 | 1.6E-03 | 9.9E-03 |
| | | | | | hsa-miR-671-5p | 2.4 | 0.6 | 2.5E-05 | 2.6E-04 |
| | | | | | hsa-miR-183-5p | 2.1 | 0.8 | 7.2E-03 | 3.6E-02 |
| | | | | | hsa-miR-1307-3p | 2.0 | 0.5 | 1.1E-04 | 9.9E-04 |
| | | | | | hsa-miR-148a-3p | 2.0 | 0.6 | 3.9E-04 | 3.0E-03 |
| | | | | | hsa-miR-574-3p | 2.0 | 0.6 | 3.1E-04 | 2.4E-03 |
| | | | | | hsa-miR-122-5p | 2.0 | 0.8 | 8.9E-04 | 6.0E-03 |
| | | | | | hsa-miR-941 | 1.7 | 0.6 | 4.0E-03 | 2.1E-02 |
| | | | | | hsa-miR-99b-3p | 1.7 | 0.6 | 3.0E-03 | 1.7E-02 |
| | | | | | hsa-miR-25-3p | 1.6 | 0.4 | 1.6E-04 | 1.4E-03 |
| | | | | | hsa-miR-185-5p | 1.5 | 0.5 | 7.1E-04 | 4.9E-03 |
| | | | | | hsa-miR-221-5p | 1.5 | 0.4 | 1.2E-03 | 8.0E-03 |
| | | | | | hsa-miR-99a-5p | -1.5 | 0.4 | 6.8E-05 | 6.1E-04 |
| | | | | | hsa-miR-100-5p | -1.6 | 0.6 | 8.4E-03 | 4.1E-02 |
| | | | | | hsa-miR-22-3p | -1.7 | 0.4 | 6.2E-06 | 8.5E-05 |
| | | | | | hsa-miR-30a-5p | -1.8 | 0.6 | 3.3E-03 | 1.7E-02 |
| | | | | | hsa-miR-21-5p | -1.9 | 0.3 | 1.9E-08 | 4.9E-07 |
| | | | | | hsa-miR-654-3p | -2.2 | 0.5 | 2.2E-05 | 2.3E-04 |
| | | | | | hsa-miR-181a-5p | -2.3 | 0.5 | 2.6E-06 | 3.7E-05 |
| | | | | | hsa-miR-29a-3p | -2.3 | 0.4 | 1.8E-08 | 4.9E-07 |
| | | | | | hsa-miR-381-3p | -2.5 | 0.6 | 1.6E-05 | 1.8E-04 |
| | | | | | hsa-miR-21-3p | -2.6 | 0.7 | 3.8E-04 | 3.0E-03 |
| | | | | | hsa-miR-199a-5p | -2.8 | 0.4 | 1.5E-12 | 9.4E-11 |
| | | | | | hsa-miR-193b-3p | -3.2 | 1.0 | 2.5E-03 | 1.4E-02 |
| | | | | | hsa-miR-136-3p | -3.5 | 0.8 | 2.5E-05 | 2.6E-04 |

**Supplementary Table 2. Differentially expressed miRNAs in Non-EVs *vs* Cells.**

| non-EV *versus* cells | | | | | | | | | |
|---|---|---|---|---|---|---|---|---|---|
| DE miRNA name | HEK | | | | DE miRNA name | MSC | | | |
| | log2FC | lfcSE | pvalue | padj | | log2FC | lfcSE | pvalue | padj |
| hsa-miR-4532 | 8.7 | 1.3 | 8.01E-10 | 2.88E-07 | hsa-miR-122-5p | 8.8 | 0.8 | 1.64E-28 | 5.69E-26 |
| hsa-miR-133a-3p | 7.3 | 1.4 | 2.51E-07 | 2.26E-05 | hsa-miR-451a | 5.5 | 1.1 | 2.33E-06 | 4.74E-05 |
| hsa-miR-3648 | 7.1 | 1.3 | 1.50E-08 | 2.70E-06 | hsa-miR-1246 | 5.4 | 1.1 | 7.44E-07 | 1.84E-05 |
| hsa-miR-4792 | 6.4 | 1.2 | 9.79E-08 | 1.18E-05 | hsa-miR-6892-5p | 5.4 | 0.9 | 2.73E-08 | 1.05E-06 |
| hsa-miR-134-5p | 6.3 | 1.4 | 1.74E-05 | 5.69E-04 | hsa-miR-6858-5p | 5.0 | 1.5 | 1.11E-03 | 8.19E-03 |
| hsa-miR-122-5p | 5.9 | 1.3 | 4.20E-06 | 2.16E-04 | hsa-miR-148a-3p | 4.9 | 0.6 | 9.71E-15 | 8.67E-13 |
| hsa-miR-663b | 5.8 | 1.5 | 7.49E-05 | 1.80E-03 | hsa-miR-760 | 4.8 | 1.0 | 7.81E-06 | 1.29E-04 |
| hsa-miR-654-5p | 5.8 | 1.5 | 1.23E-04 | 2.47E-03 | hsa-miR-365a-5p | 4.3 | 0.8 | 8.07E-08 | 2.79E-06 |
| hsa-miR-4516 | 5.7 | 1.4 | 6.63E-05 | 1.80E-03 | hsa-miR-4669 | 4.2 | 1.3 | 1.45E-03 | 9.67E-03 |
| hsa-miR-381-3p | 5.1 | 1.1 | 2.53E-06 | 1.52E-04 | hsa-miR-378a-3p | 4.2 | 1.0 | 3.96E-05 | 5.71E-04 |
| hsa-miR-432-5p | 5.1 | 1.6 | 5.71E-04 | 9.78E-03 | hsa-miR-139-5p | 4.1 | 1.3 | 1.33E-03 | 9.13E-03 |
| hsa-miR-206 | 4.9 | 1.1 | 8.70E-06 | 3.48E-04 | hsa-miR-541-5p | 4.0 | 1.4 | 4.64E-03 | 2.35E-02 |
| hsa-miR-145-3p | 4.8 | 1.7 | 4.05E-03 | 4.70E-02 | hsa-miR-150-5p | 3.7 | 1.5 | 6.25E-03 | 3.00E-02 |
| hsa-miR-143-3p | 4.8 | 1.0 | 2.20E-06 | 1.52E-04 | hsa-miR-1307-5p | 3.5 | 0.7 | 4.21E-07 | 1.12E-05 |
| hsa-miR-127-3p | 4.7 | 1.2 | 7.40E-05 | 1.80E-03 | hsa-miR-193b-5p | 3.5 | 0.7 | 3.58E-06 | 6.89E-05 |
| hsa-miR-382-5p | 4.6 | 1.4 | 1.59E-03 | 2.48E-02 | hsa-miR-378i | 3.4 | 1.2 | 3.56E-03 | 1.98E-02 |
| hsa-miR-199a-3p | 4.4 | 1.0 | 8.10E-06 | 3.48E-04 | hsa-miR-183-5p | 3.3 | 0.9 | 3.13E-04 | 2.85E-03 |
| hsa-miR-4508 | 4.3 | 1.4 | 2.80E-03 | 3.88E-02 | hsa-miR-378a-5p | 3.2 | 1.2 | 8.18E-03 | 3.63E-02 |
| hsa-miR-199b-3p | 4.3 | 1.0 | 1.12E-05 | 4.04E-04 | hsa-miR-412-5p | 3.2 | 0.8 | 7.73E-05 | 8.36E-04 |
| hsa-miR-411-5p | 4.0 | 1.3 | 2.47E-03 | 3.55E-02 | hsa-miR-6807-5p | 3.2 | 1.3 | 9.96E-03 | 4.15E-02 |
| hsa-miR-100-5p | 3.9 | 1.0 | 1.18E-04 | 2.47E-03 | hsa-miR-486-3p | 3.2 | 0.9 | 2.37E-04 | 2.28E-03 |
| hsa-miR-199a-5p | 3.5 | 0.9 | 1.85E-04 | 3.50E-03 | hsa-miR-129-5p | 3.1 | 0.9 | 2.95E-04 | 2.76E-03 |
| hsa-miR-214-3p | 3.5 | 1.2 | 3.91E-03 | 4.70E-02 | hsa-miR-320d | 3.1 | 1.1 | 6.42E-03 | 3.04E-02 |
| hsa-miR-23a-3p | 3.4 | 1.2 | 3.24E-03 | 4.17E-02 | hsa-miR-365b-5p | 3.0 | 0.7 | 5.95E-06 | 1.08E-04 |
| hsa-let-7b-5p | 3.2 | 0.9 | 4.05E-04 | 7.29E-03 | hsa-miR-204-3p | 2.9 | 1.1 | 9.48E-03 | 4.05E-02 |
| hsa-miR-22-3p | 2.5 | 0.8 | 3.13E-03 | 4.17E-02 | hsa-miR-1304-3p | 2.9 | 0.9 | 1.64E-03 | 1.03E-02 |
| hsa-miR-103a-3p | -2.5 | 0.9 | 3.84E-03 | 4.70E-02 | hsa-miR-431-3p | 2.7 | 0.9 | 3.93E-03 | 2.12E-02 |
| hsa-miR-128-3p | -2.9 | 0.9 | 1.90E-03 | 2.85E-02 | hsa-miR-134-5p | 2.7 | 0.7 | 1.86E-04 | 1.84E-03 |
| hsa-miR-615-3p | -4.6 | 1.1 | 5.04E-05 | 1.51E-03 | hsa-miR-486-5p | 2.6 | 0.8 | 1.56E-03 | 1.00E-02 |
| hsa-miR-4521 | -4.6 | 1.2 | 1.03E-04 | 2.32E-03 | hsa-miR-432-5p | 2.6 | 0.6 | 7.28E-05 | 8.12E-04 |
| hsa-miR-130b-5p | -4.7 | 1.4 | 1.24E-03 | 2.02E-02 | hsa-miR-3158-3p | 2.6 | 0.8 | 2.66E-03 | 1.56E-02 |
| | | | | | hsa-miR-23a-5p | 2.5 | 0.9 | 6.98E-03 | 3.14E-02 |
| | | | | | hsa-miR-140-3p | 2.4 | 0.6 | 7.19E-05 | 8.12E-04 |
| | | | | | hsa-miR-320c | 2.2 | 0.7 | 3.24E-03 | 1.84E-02 |
| | | | | | hsa-miR-425-3p | 2.2 | 0.7 | 1.75E-03 | 1.08E-02 |
| | | | | | hsa-miR-19b-3p | 2.2 | 0.8 | 4.22E-03 | 2.23E-02 |
| | | | | | hsa-miR-106b-3p | 2.1 | 0.7 | 2.20E-03 | 1.33E-02 |
| | | | | | hsa-miR-501-3p | 2.1 | 0.8 | 1.04E-02 | 4.28E-02 |
| | | | | | hsa-miR-192-5p | 2.0 | 0.6 | 1.53E-03 | 9.98E-03 |
| | | | | | hsa-miR-500a-3p | 1.9 | 0.8 | 1.07E-02 | 4.34E-02 |
| | | | | | hsa-miR-125a-3p | 1.9 | 0.7 | 4.25E-03 | 2.23E-02 |
| | | | | | hsa-miR-625-3p | 1.9 | 0.7 | 1.17E-02 | 4.70E-02 |
| | | | | | hsa-miR-22-5p | 1.8 | 0.7 | 6.70E-03 | 3.07E-02 |
| | | | | | hsa-miR-24-3p | 1.8 | 0.5 | 1.08E-04 | 1.14E-03 |
| | | | | | hsa-miR-25-3p | 1.8 | 0.4 | 6.57E-05 | 7.97E-04 |
| | | | | | hsa-miR-185-5p | 1.6 | 0.5 | 6.68E-04 | 5.50E-03 |
| | | | | | hsa-miR-574-3p | 1.6 | 0.6 | 5.96E-03 | 2.91E-02 |
| | | | | | hsa-miR-193a-5p | 1.5 | 0.6 | 6.63E-03 | 3.07E-02 |
| | | | | | hsa-miR-218-5p | 1.5 | 0.4 | 1.53E-04 | 1.55E-03 |
| | | | | | hsa-miR-21-5p | -1.5 | 0.3 | 1.41E-05 | 2.12E-04 |
| | | | | | hsa-miR-423-3p | -1.5 | 0.4 | 5.13E-04 | 4.33E-03 |
| | | | | | hsa-miR-199a-5p | -1.6 | 0.4 | 6.68E-05 | 7.97E-04 |
| | | | | | hsa-miR-99b-5p | -1.6 | 0.5 | 9.48E-04 | 7.29E-03 |
| | | | | | hsa-miR-127-3p | -1.6 | 0.6 | 5.13E-03 | 2.54E-02 |
| | | | | | hsa-miR-379-5p | -1.6 | 0.5 | 3.28E-04 | 2.91E-03 |
| | | | | | hsa-miR-654-3p | -1.7 | 0.5 | 1.35E-03 | 9.13E-03 |
| | | | | | hsa-miR-191-5p | -1.8 | 0.6 | 1.28E-03 | 9.13E-03 |
| | | | | | hsa-miR-487b-3p | -1.9 | 0.8 | 1.24E-02 | 4.93E-02 |
| | | | | | hsa-miR-494-3p | -2.0 | 0.6 | 8.61E-04 | 6.93E-03 |
| | | | | | hsa-miR-31-5p | -2.1 | 0.5 | 4.48E-05 | 5.96E-04 |
| | | | | | hsa-miR-221-5p | -2.1 | 0.5 | 4.19E-05 | 5.79E-04 |
| | | | | | hsa-miR-155-5p | -2.2 | 0.7 | 3.20E-03 | 1.84E-02 |
| | | | | | hsa-miR-28-5p | -2.2 | 0.8 | 4.68E-03 | 2.35E-02 |
| | | | | | hsa-miR-23b-3p | -2.4 | 0.7 | 9.32E-04 | 7.29E-03 |
| | | | | | hsa-miR-424-3p | -2.4 | 0.7 | 1.34E-03 | 9.13E-03 |
| | | | | | hsa-miR-98-5p | -2.6 | 0.8 | 1.04E-03 | 7.82E-03 |
| | | | | | hsa-miR-30c-2-3p | -2.6 | 0.9 | 2.24E-03 | 1.33E-02 |
| | | | | | hsa-miR-484 | -2.6 | 0.7 | 6.63E-05 | 7.97E-04 |
| | | | | | hsa-miR-505-3p | -2.6 | 0.9 | 3.60E-03 | 1.98E-02 |
| | | | | | hsa-miR-125a-5p | -2.6 | 0.6 | 7.11E-06 | 1.23E-04 |
| | | | | | hsa-miR-374b-5p | -2.7 | 1.0 | 6.74E-03 | 3.07E-02 |
| | | | | | hsa-miR-103a-3p | -2.9 | 0.5 | 4.33E-10 | 2.14E-08 |
| | | | | | hsa-miR-7641 | -3.0 | 1.2 | 9.10E-03 | 3.93E-02 |
| | | | | | hsa-miR-125b-5p | -3.1 | 0.5 | 5.27E-11 | 3.65E-09 |
| | | | | | hsa-miR-222-3p | -3.3 | 0.4 | 1.00E-14 | 8.67E-13 |
| | | | | | hsa-miR-671-3p | -3.9 | 0.8 | 9.37E-07 | 2.16E-05 |
| | | | | | hsa-miR-130b-5p | -4.1 | 0.9 | 1.15E-06 | 2.50E-05 |
| | | | | | hsa-let-7a-5p | -4.1 | 0.6 | 1.54E-10 | 8.89E-09 |
| | | | | | hsa-miR-455-3p | -4.2 | 1.0 | 8.41E-06 | 1.32E-04 |
| | | | | | hsa-let-7e-5p | -4.3 | 0.8 | 1.81E-07 | 5.22E-06 |
| | | | | | hsa-miR-411-5p | -4.3 | 0.5 | 5.98E-18 | 1.03E-15 |
| | | | | | hsa-miR-335-3p | -4.7 | 0.9 | 1.19E-07 | 3.75E-06 |
| | | | | | hsa-miR-411-3p | -5.3 | 0.9 | 3.23E-09 | 1.40E-07 |

**Supplementary Table 3. Differentially expressed miRNAs in EV *vs* Non-EV samples.**

| EV *versus* non-EV (part 1) | | | | | | | | | |
|---|---|---|---|---|---|---|---|---|---|
| DE miRNA name | HEK | | | | DE miRNA name | MSC | | | |
| | log2FC | lfcSE | pvalue | padj | | log2FC | lfcSE | pvalue | padj |
| hsa-miR-7704 | 8.2 | 1.2 | 3.24E-11 | 1.43E-08 | hsa-miR-1275 | 6.5 | 1.0 | 3.63E-11 | 2.80E-09 |
| hsa-miR-6087 | 7.0 | 1.1 | 1.16E-09 | 2.56E-07 | hsa-miR-128-1-5p | 6.2 | 0.9 | 2.17E-11 | 2.34E-09 |
| | | | | | hsa-miR-7977 | 5.9 | 0.9 | 1.40E-10 | 6.76E-09 |
| | | | | | hsa-miR-503-3p | 5.5 | 1.1 | 8.62E-07 | 1.75E-05 |
| | | | | | hsa-miR-671-3p | 5.4 | 0.8 | 4.42E-11 | 2.84E-09 |
| | | | | | hsa-miR-370-5p | 5.3 | 0.9 | 2.39E-08 | 7.08E-07 |
| | | | | | hsa-miR-152-5p | 5.3 | 0.9 | 8.19E-10 | 3.16E-08 |
| | | | | | hsa-miR-5100 | 5.0 | 1.1 | 4.10E-06 | 6.89E-05 |
| | | | | | hsa-miR-130b-5p | 5.0 | 0.9 | 9.34E-09 | 3.00E-07 |
| | | | | | hsa-miR-487b-5p | 4.5 | 1.2 | 5.20E-05 | 4.67E-04 |
| | | | | | hsa-miR-335-3p | 4.4 | 0.9 | 1.13E-06 | 2.07E-05 |
| | | | | | hsa-miR-323a-5p | 4.4 | 1.0 | 1.25E-05 | 1.70E-04 |
| | | | | | hsa-miR-148b-5p | 4.4 | 1.1 | 3.11E-05 | 3.16E-04 |
| | | | | | hsa-miR-23b-5p | 4.2 | 1.0 | 4.37E-05 | 4.11E-04 |
| | | | | | hsa-miR-329-5p | 4.2 | 1.3 | 7.26E-04 | 4.06E-03 |
| | | | | | hsa-miR-1225-3p | 4.0 | 1.2 | 6.05E-04 | 3.60E-03 |
| | | | | | hsa-miR-1343-3p | 3.9 | 1.0 | 8.92E-05 | 7.49E-04 |
| | | | | | hsa-miR-6789-3p | 3.9 | 1.3 | 1.90E-03 | 8.81E-03 |
| | | | | | hsa-miR-615-3p | 3.9 | 0.9 | 2.49E-05 | 2.82E-04 |
| | | | | | hsa-miR-92a-1-5p | 3.8 | 0.9 | 2.20E-05 | 2.74E-04 |
| | | | | | hsa-miR-6892-3p | 3.7 | 1.4 | 6.00E-03 | 2.17E-02 |
| | | | | | hsa-miR-410-5p | 3.7 | 1.2 | 1.88E-03 | 8.81E-03 |
| | | | | | hsa-miR-149-3p | 3.6 | 1.2 | 2.34E-03 | 1.04E-02 |
| | | | | | hsa-miR-221-5p | 3.6 | 0.5 | 5.51E-12 | 1.06E-09 |
| | | | | | hsa-miR-411-3p | 3.6 | 1.0 | 1.17E-04 | 9.39E-04 |
| | | | | | hsa-miR-29b-1-5p | 3.6 | 1.1 | 1.17E-03 | 6.01E-03 |
| | | | | | hsa-miR-92b-5p | 3.5 | 0.6 | 3.22E-08 | 8.29E-07 |
| | | | | | hsa-miR-30c-2-3p | 3.5 | 0.9 | 5.56E-05 | 4.88E-04 |
| | | | | | hsa-miR-30b-3p | 3.5 | 1.2 | 3.98E-03 | 1.62E-02 |
| | | | | | hsa-miR-4443 | 3.5 | 1.0 | 6.06E-04 | 3.60E-03 |
| | | | | | hsa-miR-25-5p | 3.5 | 0.8 | 2.70E-05 | 2.98E-04 |
| | | | | | hsa-miR-671-5p | 3.4 | 0.7 | 2.60E-07 | 5.57E-06 |
| | | | | | hsa-let-7a-5p | 3.4 | 0.6 | 1.23E-07 | 2.98E-06 |
| | | | | | hsa-miR-6807-5p | 3.3 | 1.0 | 1.74E-03 | 8.41E-03 |
| | | | | | hsa-miR-6724-5p | 3.3 | 1.2 | 4.57E-03 | 1.78E-02 |
| | | | | | hsa-miR-433-5p | 3.3 | 1.2 | 8.37E-03 | 2.80E-02 |
| | | | | | hsa-miR-455-3p | 3.2 | 1.0 | 1.04E-03 | 5.57E-03 |
| | | | | | hsa-miR-27b-5p | 3.1 | 1.1 | 4.92E-03 | 1.89E-02 |
| | | | | | hsa-let-7e-5p | 3.0 | 0.8 | 1.80E-04 | 1.29E-03 |
| | | | | | hsa-miR-33b-3p | 3.0 | 1.2 | 6.93E-03 | 2.39E-02 |
| | | | | | hsa-miR-1908-5p | 3.0 | 0.8 | 1.77E-04 | 1.29E-03 |
| | | | | | hsa-miR-411-5p | 2.9 | 0.5 | 6.10E-09 | 2.14E-07 |
| | | | | | hsa-miR-485-3p | 2.9 | 0.9 | 6.31E-04 | 3.63E-03 |
| | | | | | hsa-miR-3187-3p | 2.9 | 1.0 | 3.90E-03 | 1.60E-02 |
| | | | | | hsa-let-7a-2-3p | 2.9 | 0.9 | 2.03E-03 | 9.16E-03 |
| | | | | | hsa-miR-365b-5p | 2.8 | 0.6 | 1.28E-05 | 1.70E-04 |
| | | | | | hsa-miR-23a-5p | 2.7 | 0.8 | 1.32E-03 | 6.51E-03 |
| | | | | | hsa-miR-484 | 2.7 | 0.7 | 5.92E-05 | 5.07E-04 |
| | | | | | hsa-miR-222-3p | 2.7 | 0.4 | 1.85E-10 | 7.92E-09 |
| | | | | | hsa-miR-485-5p | 2.7 | 0.7 | 1.07E-04 | 8.77E-04 |
| | | | | | hsa-miR-409-5p | 2.7 | 0.7 | 2.96E-04 | 1.97E-03 |
| | | | | | hsa-miR-142-5p | 2.6 | 1.0 | 1.06E-02 | 3.43E-02 |
| | | | | | hsa-miR-193b-5p | 2.6 | 0.7 | 1.49E-04 | 1.13E-03 |

**Supplementary Table 3** (continued)

| EV versus non-EV (part 2) | | | | | | | | | |
|---|---|---|---|---|---|---|---|---|---|
| DE miRNA name | HEK | | | | DE miRNA name | MSC | | | |
| | log2FC | lfcSE | pvalue | padj | | log2FC | lfcSE | pvalue | padj |
| | | | | | hsa-miR-1301-3p | 2.6 | 1.1 | 1.57E-02 | 4.78E-02 |
| | | | | | hsa-miR-365a-5p | 2.5 | 0.7 | 6.43E-04 | 3.65E-03 |
| | | | | | hsa-miR-31-5p | 2.4 | 0.5 | 4.50E-06 | 7.24E-05 |
| | | | | | hsa-miR-6892-5p | 2.4 | 0.6 | 1.98E-04 | 1.39E-03 |
| | | | | | hsa-miR-98-5p | 2.4 | 0.8 | 2.80E-03 | 1.20E-02 |
| | | | | | hsa-miR-654-5p | 2.4 | 0.5 | 2.17E-06 | 3.81E-05 |
| | | | | | hsa-miR-196b-5p | 2.3 | 0.7 | 7.99E-04 | 4.41E-03 |
| | | | | | hsa-miR-193a-5p | 2.3 | 0.6 | 4.97E-05 | 4.56E-04 |
| | | | | | hsa-miR-155-5p | 2.3 | 0.8 | 3.10E-03 | 1.32E-02 |
| | | | | | hsa-miR-1307-5p | 2.2 | 0.6 | 5.30E-04 | 3.24E-03 |
| | | | | | hsa-miR-92b-3p | 2.1 | 0.6 | 9.81E-04 | 5.33E-03 |
| | | | | | hsa-miR-1307-3p | 2.1 | 0.5 | 1.37E-04 | 1.06E-03 |
| | | | | | hsa-miR-92a-3p | 2.1 | 0.5 | 1.61E-04 | 1.19E-03 |
| | | | | | hsa-miR-103a-3p | 2.1 | 0.5 | 9.01E-06 | 1.34E-04 |
| | | | | | hsa-miR-191-5p | 2.0 | 0.6 | 3.45E-04 | 2.22E-03 |
| | | | | | hsa-miR-125b-5p | 2.0 | 0.5 | 2.45E-05 | 2.82E-04 |
| | | | | | hsa-miR-424-3p | 1.9 | 0.8 | 1.24E-02 | 3.87E-02 |
| | | | | | hsa-miR-23b-3p | 1.9 | 0.7 | 8.01E-03 | 2.71E-02 |
| | | | | | hsa-miR-125a-5p | 1.9 | 0.6 | 1.30E-03 | 6.51E-03 |
| | | | | | hsa-miR-24-2-5p | 1.8 | 0.7 | 1.55E-02 | 4.74E-02 |
| | | | | | hsa-miR-328-3p | 1.8 | 0.7 | 1.38E-02 | 4.25E-02 |
| | | | | | hsa-miR-423-3p | 1.6 | 0.4 | 2.29E-04 | 1.55E-03 |
| | | | | | hsa-miR-196a-5p | 1.6 | 0.6 | 6.13E-03 | 2.17E-02 |
| | | | | | hsa-miR-433-3p | 1.6 | 0.6 | 1.14E-02 | 3.60E-02 |
| | | | | | hsa-miR-181b-5p | -1.6 | 0.5 | 4.95E-04 | 3.13E-03 |
| | | | | | hsa-miR-99a-5p | -1.6 | 0.4 | 2.90E-05 | 3.02E-04 |
| | | | | | hsa-miR-532-5p | -1.7 | 0.6 | 2.56E-03 | 1.12E-02 |
| | | | | | hsa-miR-30e-5p | -1.8 | 0.7 | 1.11E-02 | 3.59E-02 |
| | | | | | hsa-miR-767-5p | -1.8 | 0.6 | 5.32E-03 | 2.01E-02 |
| | | | | | hsa-miR-339-3p | -1.8 | 0.6 | 3.33E-03 | 1.38E-02 |
| | | | | | hsa-miR-218-5p | -1.8 | 0.4 | 8.68E-06 | 1.34E-04 |
| | | | | | hsa-miR-30d-5p | -1.9 | 0.7 | 6.62E-03 | 2.30E-02 |
| | | | | | hsa-miR-143-3p | -1.9 | 0.5 | 3.10E-04 | 2.03E-03 |
| | | | | | hsa-miR-22-3p | -2.0 | 0.4 | 1.95E-07 | 4.43E-06 |
| | | | | | hsa-miR-134-5p | -2.0 | 0.7 | 4.57E-03 | 1.78E-02 |
| | | | | | hsa-miR-24-3p | -2.1 | 0.5 | 1.02E-05 | 1.46E-04 |
| | | | | | hsa-miR-425-3p | -2.1 | 0.8 | 8.49E-03 | 2.80E-02 |
| | | | | | hsa-miR-101-3p | -2.1 | 0.9 | 1.22E-02 | 3.84E-02 |
| | | | | | hsa-miR-432-5p | -2.1 | 0.7 | 1.25E-03 | 6.35E-03 |
| | | | | | hsa-miR-186-5p | -2.2 | 0.7 | 1.12E-03 | 5.90E-03 |
| | | | | | hsa-miR-181a-5p | -2.2 | 0.5 | 1.83E-05 | 2.36E-04 |
| | | | | | hsa-miR-320c | -2.3 | 0.8 | 5.44E-03 | 2.04E-02 |
| | | | | | hsa-miR-105-5p | -2.4 | 0.9 | 6.55E-03 | 2.30E-02 |
| | | | | | hsa-miR-382-3p | -2.4 | 0.8 | 1.80E-03 | 8.58E-03 |
| | | | | | hsa-miR-22-5p | -2.5 | 0.8 | 1.14E-03 | 5.94E-03 |
| | | | | | hsa-miR-148a-3p | -2.6 | 0.6 | 2.85E-05 | 3.02E-04 |
| | | | | | hsa-miR-136-5p | -2.6 | 0.9 | 4.49E-03 | 1.78E-02 |
| | | | | | hsa-miR-148b-3p | -2.7 | 0.6 | 2.44E-05 | 2.82E-04 |
| | | | | | hsa-miR-192-5p | -2.7 | 0.7 | 1.29E-04 | 1.01E-03 |
| | | | | | hsa-miR-29a-3p | -2.7 | 0.4 | 8.90E-11 | 4.91E-09 |
| | | | | | hsa-miR-27a-3p | -2.8 | 0.4 | 2.43E-11 | 2.34E-09 |
| | | | | | hsa-miR-410-3p | -2.8 | 1.0 | 7.66E-03 | 2.62E-02 |
| | | | | | hsa-miR-345-5p | -2.9 | 1.0 | 5.92E-03 | 2.17E-02 |
| | | | | | hsa-miR-376c-3p | -2.9 | 1.1 | 1.12E-02 | 3.59E-02 |
| | | | | | hsa-miR-30a-5p | -3.0 | 0.6 | 1.04E-06 | 2.00E-05 |
| | | | | | hsa-miR-21-3p | -3.2 | 0.8 | 4.17E-05 | 4.02E-04 |
| | | | | | hsa-miR-381-3p | -3.2 | 0.6 | 2.68E-08 | 7.39E-07 |
| | | | | | hsa-miR-193b-3p | -3.3 | 1.1 | 3.16E-03 | 1.33E-02 |
| | | | | | hsa-miR-1246 | -3.4 | 1.1 | 2.04E-03 | 9.16E-03 |
| | | | | | hsa-miR-660-5p | -3.5 | 1.1 | 4.23E-03 | 1.70E-02 |
| | | | | | hsa-miR-136-3p | -3.5 | 0.8 | 3.99E-05 | 3.94E-04 |
| | | | | | hsa-miR-376a-3p | -3.5 | 1.2 | 5.76E-03 | 2.14E-02 |
| | | | | | hsa-miR-424-5p | -3.7 | 1.4 | 8.45E-03 | 2.80E-02 |
| | | | | | hsa-miR-377-3p | -4.0 | 1.2 | 1.47E-03 | 7.18E-03 |
| | | | | | hsa-miR-548o-3p | -4.1 | 1.3 | 1.92E-03 | 8.81E-03 |
| | | | | | hsa-miR-19b-3p | -4.1 | 1.0 | 2.15E-04 | 1.48E-03 |
| | | | | | hsa-miR-760 | -4.6 | 1.3 | 6.16E-04 | 3.60E-03 |
| | | | | | hsa-miR-122-5p | -6.2 | 0.8 | 6.80E-15 | 2.62E-12 |
